# Flexible communication between cell assemblies and ‘reader’ neurons

**DOI:** 10.1101/2022.09.06.506754

**Authors:** Marco N. Pompili, Ralitsa Todorova, Céline J. Boucly, Eulalie M. Leroux, Sidney I. Wiener, Michaël Zugaro

## Abstract

Cell assemblies are considered fundamental units of brain activity, underlying diverse functions ranging from perception to memory and decision-making. Cell assemblies have generally been studied in relation to specific stimuli or actions, but this approach does not readily extend to more abstract constructs. An alternative approach is to assess cell assemblies without making reference to external variables, and instead focus on internal brain processes — by defining assemblies by their endogenous ability to effectively elicit specific responses in downstream (‘reader’) neurons. However, this compelling idea currently lacks experimental support. Here, we provide evidence for assembly–reader communication. Reader activation was genuinely collective, functionally selective, yet flexible, implementing both pattern separation and completion. These processes occurred at the time scale of membrane integration, synaptic plasticity and gamma oscillations. Finally, assembly–reader couplings were selectively modified upon associative learning, indicating that they were plastic and could become bound to behaviorally relevant variables. These results support cell assemblies as an endogenous mechanism for brain function.

---

An increasingly influential hypothesis in neuroscience posits that the basic unit of brain processing consists of ‘cell assemblies’ (*1–6*), a term coined by Hebb to designate groups of neurons that were hypothesized to activate during specific cognitive processes. Although the precise structure and timescale of cell assemblies were not explicitly defined in this initial description, recent investigations have generally converged to the notion of a very dynamic and transient co-activation of specific neuronal ensembles, occurring in brief time windows of at most a few dozen milliseconds (*7, 8*). Such recurring patterns of coincident activity are thought to mediate complex information processing beyond the aggregate power of single cells.

Cell assemblies have generally been studied in relation to controlled stimuli or motor activity, with the underlying assumption that assembly members can code for complementary features that become bound through synchronous activation (*9–11*). However, this approach cannot be readily extended to higher order cortical areas, where assemblies (*12–20*) cannot easily be assessed in terms of known sensory or motor correlates, and where represented entities, such as psychological states (intentions, beliefs, etc.), are ill defined and notoriously difficult to test. More fundamentally, epistemological arguments suggest that this may not simply be due to contingent experimental limitations: commonly posited cognitive functions may or may not be appropriate depictions of actual brain functions (*21*).

How then should one study cell assemblies without referencing experimental parameters outside the brain, such as sensory or behavioral variables? One possibility is to instead focus on brain processes, and consider the effects of assemblies on downstream ‘reader’ neurons: in this alternative, ‘brain-based’ approach, a cell assembly is characterized by its ability to trigger a specific response in one or more target neurons, underlying specific autonomic, behavioral or cognitive functions — Buzsáki goes as far as to suggest that “the cell assembly can only be defined from the perspective of a reader mechanism” (*22*). Importantly, this also shifts the focus from a fundamentally correlational question, centered on temporal coincidences between cell assemblies and behavioral or psychological variables, to the causal question of the action of cell assemblies on downstream circuits (see (*23*)).

However, whether this appealing framework is supported by physiological evidence remains an open question, possibly because of technological limitations even in state-of-the-art causal technologies. Indeed, current approaches, including timed and targeted optogenetic manipulations, do not yet permit selective and thorough manipulation of defined groups of neurons only at the precise moment when they are about to form specific cell assemblies, but not when they fire individually. Our alternative approach to causally test this conceptual framework is to use behavioral intervention to demonstrate that assembly-reader couplings, identified during endogenous activity (sleep), can be altered in a predictable manner by behavioral paradigms designed to induce learning and memory in the respective brain structures.

## Results

### Identification of candidate cell assemblies

We recorded from large neuronal ensembles in two reciprocally interconnected associative brain areas, namely the prefrontal cortex-amygdala circuit (Fig. 1a) (*24*). In order to avoid referencing variables outside the brain, we examined collective neural dynamics during sleep, when the brain is minimally engaged in processing sensory inputs and motor outputs, and instead its activity is dominated by endogenous processes.

**Fig. 1.**
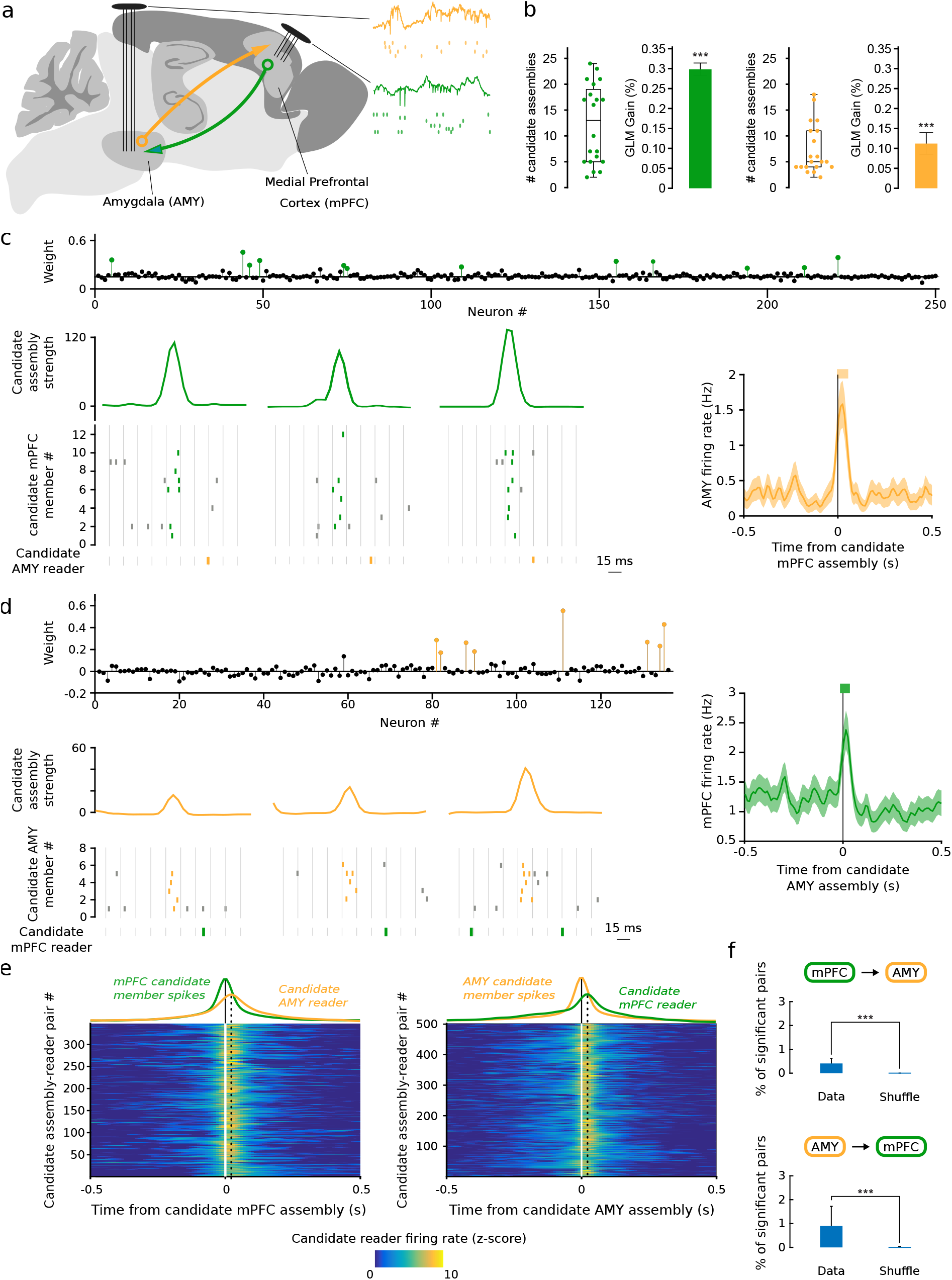
Candidate cell assembly activations are closely followed by downstream spiking. **a**, Simultaneous high-densisty recordings in the bi-directionally connected medial prefrontal cortex and amygdala (*n* = 4 rats; 5 sessions each). **b**, Numbers of candidate assemblies in prefrontal (left two panels, median *n* = 13) and amygdalar (right two panels, median *n* = 5) recordings per session (boxes and whiskers: distribution quartiles). Bars: peer prediction of the activity of candidate assembly members from other members (gain relative to shuffled data, median *±* s.e.m.; \*\*\**p <* 0.001, Wilcoxon rank sum test). **c**, Example activations of candidate medial prefrontal cortical assembly closely followed (10–30 ms) by responses of an amygdalar neuron. Top: candidate cell assembly weights (colored circles: assembly members, black circles: non-members). Bottom left: examples of candidate assembly activation (curves: activation strength) followed by downstream spiking (rasters: prefrontal spikes within (green) and outside (gray) epochs of candidate assembly activation; orange rasters: amygdalar spikes). Right: firing rate of amygdalar neuron centered on all prefrontal candidate assembly activations (mean ± s.e.m.) Thick orange horizontal bar indicates significant responses (*p <* 0.05: Monte-Carlo bootstrap test; see Methods). **d**, Same as (**c**) for candidate amygdalar assembly and downstream prefrontal neuron. **e**, Average downstream responses (z-scored firing rates) centered on candidate assembly activations, over all significant pairs (color plots), and averaged across pairs (color curves) compared with the average activity of the upstream candidate assembly. Left: candidate prefrontal assemblies and amygdalar downstream neurons. Right: candidate amygdalar assemblies and downstream prefrontal neurons. **f**, Percentage of significant candidate assembly-reader pairs found in shuffled recordings vs. real data (\*\*\**p <* 0.001, Wilcoxon signed-rank test).

The first step was to identify ensembles of coactive neurons. While ensembles characterized by pure spike train statistics are often considered in the literature to qualify as ‘cell assemblies’, given the theoretical context of the present work, we will provisionally term them ‘candidate’ cell assemblies while testing whether they are detected by downstream neurons. Indeed, a more complete characterization would require attribution of additional functional properties, such as feature coding (traditional approach) or effective spike transmission (brain-based approach tested here). Candidate cell assemblies were identified using a classical PCA-ICA algorithm (*25*) (see Methods). In each sleep session, groups of prefrontal units recurrently fired with high synchrony, forming candidate cell assemblies (*13–16*) (Fig. 1b,c; Fig. S1a). This did not spuriously arise from global fluctuations in firing rates during sleep (Fig. S1f).

We verified that candidate cell assemblies corresponded to ensembles of neurons that fired uniquely synchronous spike patterns. Indeed, cells participating in candidate assemblies (‘members’) fired more synchronously with each other than with non-members (Fig. S1c-e), and their spike trains could be reliably predicted from those of other members of the same candidate assembly (‘peer prediction’, (*12*); Fig. 1b).

Since the prefrontal-amygdalar pathway has reciprocal connections, we also tested for coactivation of amygdalar neurons. Indeed, amygdalar neurons formed candidate cell assemblies (Fig. 1b,d, Fig. S1b) (*19, 26–28*), which did not spuriously arise from global fluctuations in firing rates during sleep (Fig. S1f). Similar to the prefrontal cortex, synchrony and peer prediction were significantly greater than expected by chance (Fig. 1b, Fig. S1c-e).

### Demonstration of reader neuron responses to assembly activations

According to the brain-based framework, assembly activations should effectively elicit discharges in down-stream reader neurons. This has two implications: first, activation of an assembly should precede that of its reader within a brief time window, occurring more frequently than expected by chance; and second, this relationship should be dependent on the *collective* activation of the members of the assembly.

We first investigated whether candidate prefrontal cell assemblies reliably triggered spiking in downstream amygdalar neurons within 10–30 ms, corresponding to the conduction delay between the two structures in the rat brain (*29*). In 347 candidate assembly–reader pairs (Fig. 1e,f) this temporal coordination was greater than expected by chance (*p <* 0.05, Monte-Carlo bootstrap). Conversely, in 502 cases, candidate amygdalar assembly activations were consistently followed by prefrontal spikes (*p <* 0.05, Monte-Carlo bootstrap; Fig. 1e,f; see also Fig. S2). We confirmed that, for both prefrontal and amygdalar assembly activations, the probability that significant responses in downstream neurons peaked within the 10– 30 ms time window (*29*) (Fig. S3). Finally, consistent with the hypothesis that collective activation of cell assembly members drives responses in reader neurons, downstream neurons were more likely to discharge when increasing numbers of members were active together (Fig. S4).

Second, to assess whether spiking in downstream neurons was actually selective for the collective activation of upstream candidate assemblies, we sought to rule out two possibly confounding scenarios: 1) downstream neurons could be merely responding to each of the candidate assembly members independently, and 2) they could be responding to the compound excitatory drive exerted by active members, irrespective of how many and which different members were actually active.

We first verified that candidate assembly members exerted a synergistic, rather than independent (linearly summating), influence on their targets. In one extreme scenario, one or two particularly influential members might suffice to evoke maximal discharge in the target neuron while the other members would not impact the response at all. Thus, we reanalyzed the data using only activations of the candidate assemblies in which the members that evoked greater responses in readers remained silent (these could be identified since not all members are recruited in all activations). Even then, the responses of the target neurons remained well above their baseline firing rates, indicating that the target neuron did not only respond to a ‘vocal’ minority of the assembly members. (Fig. 2a; Fig. S5).

**Fig. 2.**
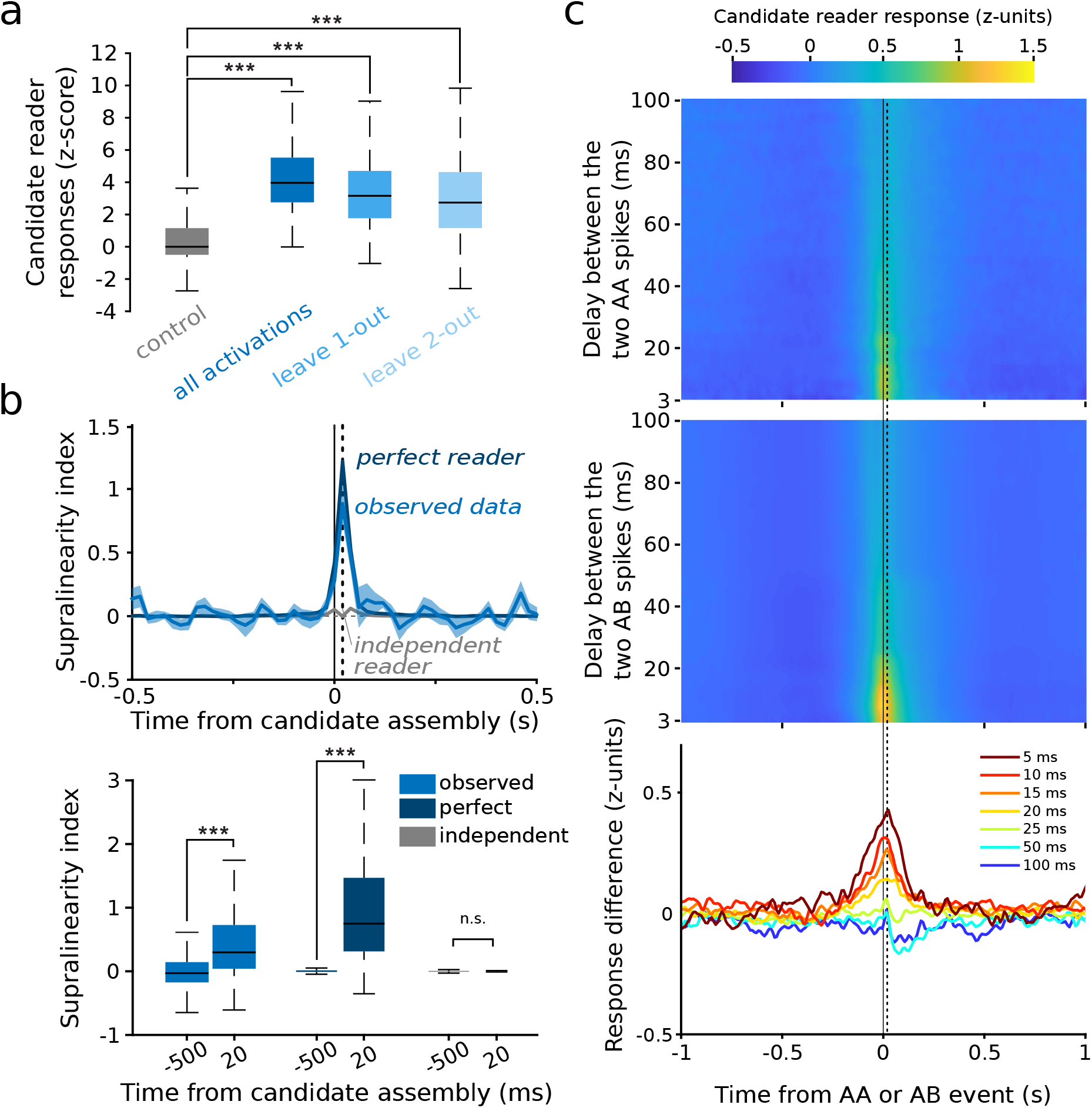
Readers respond to the collective activity of assembly members. **a**, Average responses of reader neurons to those candidate assembly activations when the most influential members of candidate assemblies were not recruited. As a control, candidate assemblies and downstream neurons were taken randomly among non-significant pairs (mPFC and AMY pooled; shown separately in Fig. S5). **b**, Supralinearity of reader responses to the collective activity of candidate assembly members. Top: Supralinearity index of data (blue curve) compared to a simulated ‘perfect’ collective reader (dark blue curve) and to a simulated ‘independent’ reader (gray curve). Bottom: The supralinearity indices 20 ms after candidate assembly activations were significantly greater than at baseline (500 ms prior to assembly activations) for both the observed data (\*\*\**p <* 0.001, Wilcoxon signed-rank test) and the simulated perfect collective readers (\*\*\**p <* 0.001, Wilcoxon signed-rank test), but not for the simulated independent readers (p=0.7916, Wilcoxon signed-rank test). **c**, Top: mean z-scored responses of reader neurons to two successive spikes of the same member (AA) of a candidate assembly for different delays between the two spikes. Center: same as top, but for reader responses to two successive spikes of two different candidate assembly members (AB). Bottom: difference between the two (AB AA), for varying temporal delays. The response to co-activations of different members (AB) is greater than the response to multiple activations of the same member (AA) only for brief (*<*25 ms) delays between spikes (\*\*\**p <* 0.001, Wilcoxon signed-rank test). Vertical dashed lines indicate 20 ms (counted from the mean of the AA or AB timestamps).

To further address this scenario in its most general form, we tested whether the reader response exceeded what could be expected from linear summation alone, i.e. whether it was supralinear. To estimate a linear summation response, we first trained a generalized linear model (GLM) to predict responses to spikes emitted by single members (i.e., at times when candidate assemblies were not active), and we then used this pre-trained GLM to predict the reader response to the activation of candidate assemblies. The observed reader response exceeded this linear summation estimate, confirming that the assembly-reader mechanism involves supralinear coincidence detection (Fig. S6). We compared the observed supralinearity to two simulated readers: a ‘perfect’ reader, which responds exclusively to the collective activity of the candidate assembly, and an ‘independent’ reader, which responds to members independently (see Methods). The observed reader responses closely resembled the responses of the simulated ‘perfect’ reader, significantly surpassing the GLM-predicted activity, and this supralinearity peaked at a delay of 20 ms Fig. 2b), indicating that the collective activation of candidate assembly members was capable of evoking greater responses than the sum of their individual contributions.

We then assessed whether readers detected the synchronous activity of specific multiple members of the candidate assemblies, or simply responded to a compound drive (total spike count) by subsamples of the same presynaptic neurons. We therefore tested if members were interchangeable, or even dispensable, provided their total spike count remained the same. To test for this, for any given pair of candidate assembly members (A and B), we compared reader responses when each of the two members emitted exactly one spike (AB) vs. when only one of the two members emitted exactly two spikes (AA), thus maintaining a constant number of candidate assembly spikes while altering which subgroup of members participated in the event. This analysis revealed that the collective activity of members mattered beyond their compound drive (Fig. 2c, Fig. S7). This is consistent with the notion that the response of the reader neuron depends on detailed spatio-temporal properties of its inputs (e.g., precisely timed spike patterns impinging on specific combinations of dendritic branches (*30*)).

We hypothesized that a given assembly could drive multiple reader neurons which may, in turn, participate in cell assemblies. Indeed, 82 of the 204 amygdalar readers participated in 147 candidate assemblies, 42 of which were, in turn, detected by prefrontal readers. Further, compared to other amygdalar neurons, amygdalar readers were significantly more likely to participate in cell assemblies targeting prefrontal readers (*p* = 1.2 10*^−^*^4^, *χ*^2^ test). Similarly, 278 prefrontal readers (out of 404) participated in 247 candidate assemblies, 104 of which triggered amygdalar reader firing, and thus were significantly more likely than other prefrontal neurons to target amygdalar readers (*p* = 2.6 10*^−^*^21^, *χ*^2^ test). Thus cell assemblies can be detected by cell assemblies, extending the concept of reader neuron,s and providing a generalized mechanism for bidirectional communication.

### Properties of the assembly-reader mechanism

These results support the prediction that assemblies exert a collective impact on their readers. To investigate the time scale of this synergistic effect, we repeated these analyses for varying interspike intervals and assembly activation durations. Both approaches yielded results consistent with an endogenous time scale of up to ∼ 20-25 ms for effective cell assemblies (Fig. 2c, Fig. S8, and Fig. S9). This time scale corresponds to those of functionally relevant cellular and network properties, including membrane integration time constants and local delays (*31*), optimal time windows for spike timing dependent potentiation of synaptic efficacy (*32*), and the period of synchronizing gamma oscillations (*33*).

We next investigated computational and functional properties of the assembly–reader mechanism, again focusing on brain dynamics, in line with the brain-based framework. Two generic computations of fundamental theoretical relevance (*37*) are pattern completion, which allows accurate retrieval or generalization from partial inputs, and pattern separation, which improves discrimination of similar inputs (*4,35,37,44*). These have been ubiquitously identified from sensory-motor to associative networks (e.g. in the cerebellum (*34*), hippocampal system (*35, 36*), cortex (*37*), olfactory networks (*38–40*), visual networks (*41, 42*), amygdala (*43*), etc.) Do readers respond similarly upon activation of a sufficient subset of assembly members, and do they discriminate between assemblies with common members? Operationally, these properties can be tested by examining relations between input and output activity patterns, which should have a sigmoid shape (*45–48*).

To test for pattern completion, we assessed reader responses following partial activation of upstream assemblies, and measured how they increased with the number of active members. Reader responses did not simply increase proportionally to the number of active members, but rather were significantly better fit by a sigmoid curve, thus providing evidence for pattern completion (*49*) (Fig. 3a-b, Fig. S10). Regarding pattern separation, we verified that reader neurons clearly discriminated between different assemblies, even when these included a number of common members (overlapping assemblies). As a first step, we compared how reader neurons responded to each of the simultaneously recorded assemblies, and found that the distributions of their responses were sparser than similar distributions constructed from non-reader activity or shuffled data (Fig. 4a, Fig. S11). This confirmed that readers were highly selective for specific assemblies. We then further verified that reader neurons could even discriminate between substantially overlapping assemblies. Indeed, reader firing rates were significantly higher following activation of their associated assembly than following activation of different assemblies with common members to their associated assembly (≥ 25% overlap) (Fig. 4b, Fig. S12). Overall, these results show that the assembly–reader mechanism is both robust and selective, since it can implement both pattern completion and pattern separation.

**Fig. 3.**
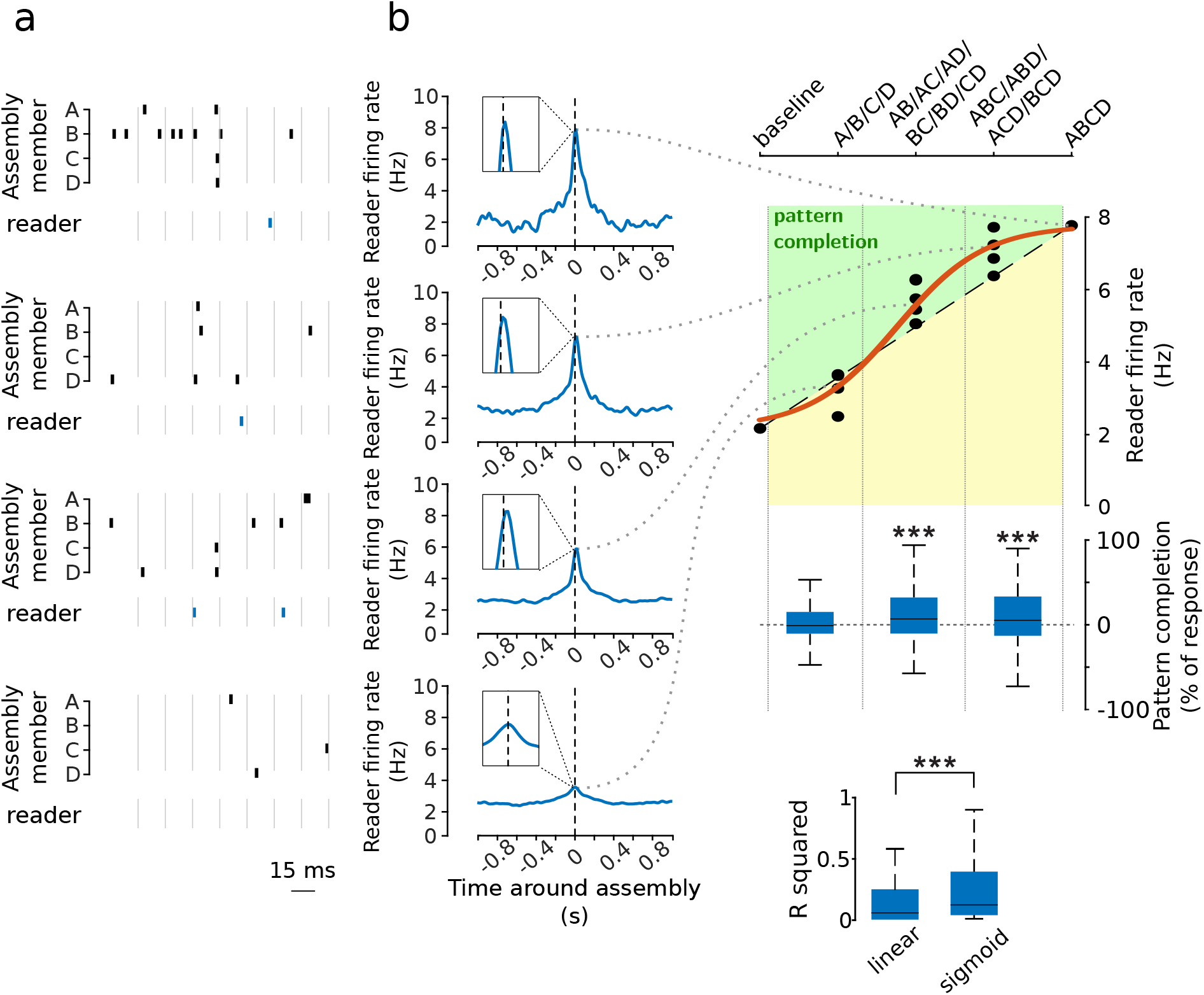
Pattern completion in assembly–reader pairs. **a** Example activity patterns of assembly members (black ticks) and a reader (blue ticks). **b**, Response of the reader in (a) to incomplete activations of a 4-member assembly (ABCD). Dashed line: proportional response. Red curve: best-fit sigmoid. Green zone: pattern completion. Center: boost in reader response (relative to a proportional response) for all assembly– reader pairs as a function of the number of active assembly members. The gain was significant for the second and third quantiles (\*\*\**p <* 0.001, Wilcoxon signed-rank test). Bottom: Proportional vs sigmoidal fits of observed data (\*\*\**p <* 0.001, Wilcoxon signed-rank test).

**Fig. 4.**
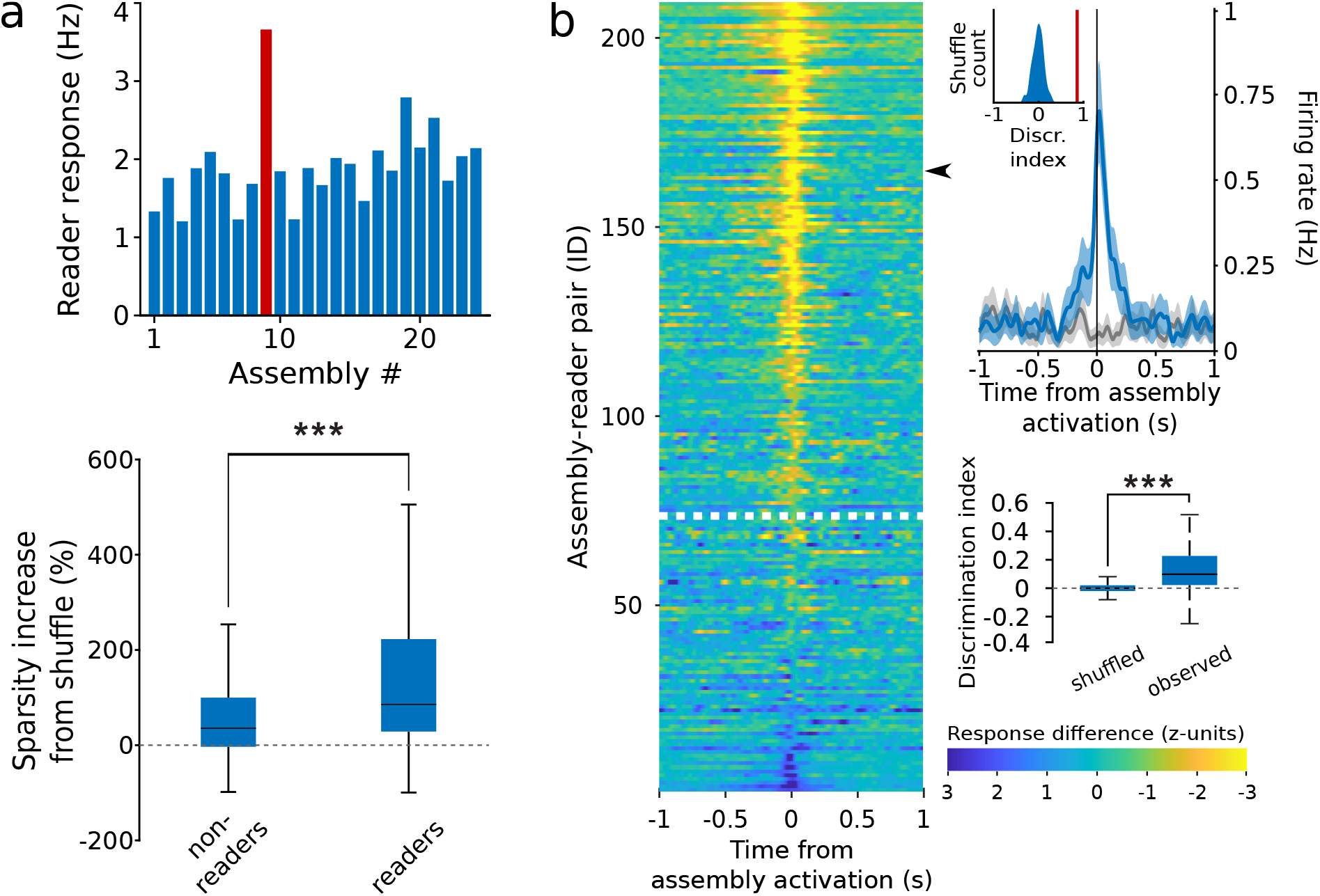
Pattern separation in assembly–reader pairs. **a**, Pattern separation. Top: responses of an example reader neuron to each cell assembly detected in the same session (red bar: assembly specific to this downstream neuron). Bottom: sparsity (increase relative to shuffled data) of the responses of reader neurons to assembly activations was significantly greater than shuffled data (*p <* 0.001, Wilcoxon signed rank test) and than responses of non-reader neurons (\*\*\**p <* 0.001, Wilcoxon rank sum test). **b**, Pattern separation. Left: Mean difference between reader responses to activations of paired assemblies and other assemblies with overlapping members (≥ 25% of all members). Data are sorted by discrimination index. Responses above the white dotted line displayed significant pattern separation (discrimination index greater than 95% of the shuffled data). Top right: response of an example reader (black arrow) to its paired assembly (blue curve) vs to another assembly with overlapping members (gray curve) (mean ± s.e.m.). Bottom right: Observed discrimination indices were greater than the discrimination indices for shuffled data (\*\*\**p <* 0.001, Wilcoxon rank sum test).

### Learning-related plasticity of the assembly-reader mechanism

Next, we tested the prediction that assembly-reader relations could be altered by learning and memory in a selective and predictable manner. We compared assembly–reader pairs before and after a standard fear conditioning and extinction protocol known to recruit the prefronto-amygdalar circuit (*50–53*) (Fig. 5a; see Methods). During fear conditioning and subsequent sleep, fear-related signals would be expected to flow from the amygdala to the prefrontal cortex (*54*), leading to consolidation of fear memories (*14,17,55*). Prefrontal reader responses to amygdalar assemblies were compared between sleep sessions preceding vs following training. Many amygdalar cell assemblies were active in both sleep sessions, but formed novel associations with downstream prefrontal neurons following fear conditioning (Fig. 5a). Conversely, other downstream prefrontal neurons no longer responded significantly to amygdalar cell assemblies in post-conditioning sleep (Fig. 5a). These results suggest that fear conditioning induced a reorganization of the functional relations between assemblies and readers, mediating the formation of conditioning-related memory traces spanning the amygdala and the prefrontal cortex.

**Fig. 5.**
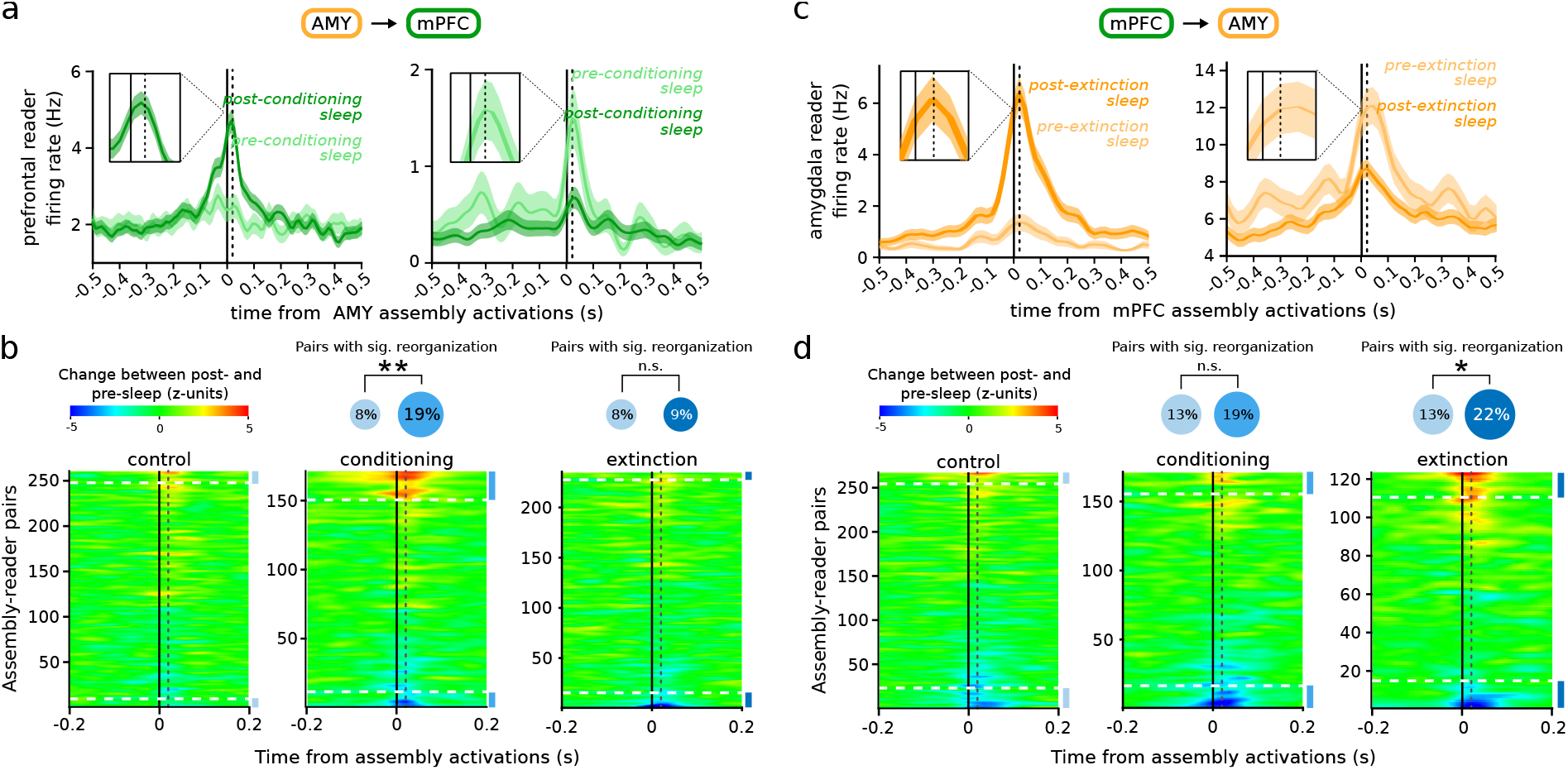
Learning-related changes in assembly–reader relations. **a**, Examples readers increasing (left) or decreasing (right) their responses (shaded area: mean ± s.e.m.) to assembly activations before (light green) and after (green) fear conditioning. Dotted line: 20 ms. **b**, Left: Differences in responses of a prefrontal reader between post- and pre-sleep, centered on amygdalar assembly activations. Data are sorted according to response magnitudes. Assembly–reader pairs above the higher dashed line (respectively, below the lower dashed line) significantly (*p <* 0.05, Monte-Carlo bootstrap test) increased (respectively, decreased) their responses in post-task sleep. A greater proportion of assembly–reader pairs significantly changed their responses (*p <* 0.05, Monte-Carlo bootstrap test) following fear conditioning (middle, blue disk) than following control sessions (middle, cyan disk; \*\**p* = 0.0017, chi-square test). In contrast, the number of assembly–reader pairs that significantly changed their responses was not greater after fear extinction (right, dark blue disk) than after control sessions (*p >* 0.05, chi-square test). **c**, Same as (**a**) for example amygdalar reader responses to prefrontal assemblies following fear extinction. **d**, Same as (**b**) for amygdalar reader responses to prefrontal assemblies. A greater proportion of assembly–reader pairs significantly changed their response (*p <* 0.05, Monte-Carlo bootstrap test) after fear extinction (right, dark blue disk) than after control sessions (right, blue disk; \**p* = 0.0347, chi-square test). In contrast, the number of assembly–reader pairs that significantly changed their responses was not greater after fear conditioning (middle, blue disk) than after control sessions (*p >* 0.05, chi-square test).

To confirm that these changes were specifically related to fear learning (as opposed to, e.g., changes in motor activity), we recorded responses before and after a control session where no conditioning took place. Compared to this control, fear conditioning was followed by significantly greater responses in prefrontal readers to amygdalar assemblies (Fig. 5b). Further, in contrast to fear conditioning, fear extinction did not result in such changes (Fig. 5a), indicating that variations in assembly–reader relations were not broadly elicited by general fearful behavior, but rather reflected the specific process of forming new fear memories.

Conversely, during fear extinction, the prefrontal cortex would be expected to alter amygdalar signals (*56*). Consistent with this, the relation between prefrontal assemblies and amygdalar readers underwent substantial reorganization following fear extinction (Fig. 5c). Again, this was specific to this particular cognitive process, since fear conditioning did not yield changes in assembly-reader pairs significantly different from control sessions (Fig. 5b).

## Discussion

In recent years, cell assemblies have become a central research topic in neuroscience. Numerous studies have investigated how cell assemblies may encode external stimuli, motor commands, or cognitive variables (*9–20*). Often implicit in this approach is the notion that the brain operates as a representational device: external stimuli are expected to trigger precise synchronous activity in dedicated ensembles of neurons, which are conceived of as internal representations of the stimuli. Yet, on the basis of epistemological, psychological, anatomical or physiological arguments, one can question this feedforward view of an essentially passive brain awaiting inputs to compute optimal outputs (*22, 57–60*). A similar argument can be made for more abstract (non-sensory) constructs, including psychological processes such as attention, volition or imagination, for which it may be illusory to seek brain mechanisms (*21*). Further, these approaches fail to address the causal mechanisms of the formation or impact of cell assemblies (for a discussion of spike causality, see e.g. (*23*)). What alternative approach could help us better understand the brain mechanisms of higher order, cognitive functions? It has been suggested that focusing on brain dynamics could provide novel insights (*60*). Thus, cell assemblies could be defined not by their mapping to external stimuli, but by their unique, causal ability to trigger specific responses in downstream neurons (*22*). However, it remained unclear that the synchronous activity of specific neuronal ensembles is uniquely relevant for downstream circuits.

Synchronous neuronal ensembles have been documented in various brain regions: in the hippocampus (*12, 20*), in the medial prefrontal cortex (*13–16*), in sensory cortices (*61*), in motor cortices (*62*), as well as in the basal ganglia (*63*) and amygdala (*28*). Such ensembles have even been reported to span multiple brain areas (*19*). These studies indicate that certain external (or internal) conditions coincide with the formation of putative cell assemblies, suggesting a mechanistic relation to behavioral or cognitive functions. However, they do not yet provide evidence that the remarkable co-activity patterns detected by the experimenters are also detected by the brain itself.

We showed that neurons in downstream structures can reliably and selectively respond to the activation of upstream co-active ensembles. According to the brain-based framework, such downstream neurons can thus be deemed ‘readers’, and upstream ensembles, ‘cell assemblies’. Indeed, reader responses were stronger than expected for the sum of independent inputs, and depended on the identity of the participating neurons rather than their aggregate drive. While supralinear responses could be expected from modeling and biophysics experiments (*66*), evidence was lacking that under physiological conditions in the living brain, post-synaptic neurons could selectively respond to specific patterns of coincident presynaptic spikes, but not to equivalent inputs originating from different combinations of neurons. In particular, our results indicate that readers respond more strongly to spikes emitted by more numerous members of a presynaptic assembly than to the same number of spikes emitted by fewer members of the same assembly. The assembly-reader coupling therefore implements a genuinely collective process, and supports a possible alternative, operational, rather than representational, definition for cell assemblies (*22*), without reference to features of external stimuli or actions (*8*).

Importantly, individual neurons were involved in both cell assembly and reader functions, suggesting that the readout of cell assemblies was not only performed by isolated neurons, but more generally by other cell assemblies as well (*67,68*). This suggests a communication scheme whereby cell assemblies transiently become dynamically coupled to other cell assemblies in the same or other, even multiple, brain areas. In addition, a fraction of the upstream assembly members were sufficient to elicit reader responses, indicating that readers could reliably respond even to incomplete assemblies, effectively implementing a pattern completion process (*4, 37, 44–46, 48, 49*). Conversely, readers discriminated between partly overlapping assemblies, indicating that they could resolve ambiguous signals by separating similar inputs, effectively implementing pattern separation (*4, 37, 45–49*). Thus, we showed that the assembly-reader mechanism is robust and selective since it implements both pattern separation and pattern completion.

The above results suggest that communication between cell assemblies can be effective and selective, and mediate advanced capabilities long hypothesized to be instrumental for complex cognitive functions. To demonstrate the functional role of such assembly-reader couplings, we subjected rats to fear conditioning and extinction protocols. Our results showed that flexible functional changes in assembly–reader pairing emerged during learning. In particular, there was a double dissociation between learning and extinction, on one side, and changes in prefrontal and amygdalar reader responses, on the other side. Previous findings indicate that fear-related signals should flow from the amygdala to the prefrontal cortex (*54*), whereas fear extinction should be mediated by prefrontal cortical control of amygdalar signals (*56*). Consistently, we found that fear conditioning elicited changes in amygdalar reading of prefrontal assemblies, whereas fear extinction elicited changes in prefrontal reading of amygdalar assemblies. This provided evidence that assembly–reader relations can selectively change during learning in a behaviorally-relevant manner, supporting the role of cell assemblies as functional units of brain dynamics, within the non-representational framework of brain function.

## Acknowledgments

We thank G. Makdah for help with data acquisition and Y. Dupraz for technical support. This project was funded by the Agence Nationale de la Recherche (ANR-17-CE37-0016-01) (M.Z.), Fondation pour la Recherche Médicale (Équipe FRM EQU202103012768) (M.Z.), Labex MemoLife (ANR-10-LABX-54 MEMO LIFE, ANR-10-IDEX-0001-02 PSL*) (M.N.P. and S.I.W.), French Ministry of Research (C.B.), and Collège de France (R.T.) The authors declare no competing interests. Research design: C.J.B., M.N.P., R.T., S.I.W., M.Z. Experiments: E.M.L., M.N.P. Data analysis design: C.J.B., M.N.P., R.T., S.I.W., M.Z. Data analysis: C.B.J., R.T. Manuscript: C.B.J., M.N.P., R.T., S.I.W., M.Z.

## Animals

Four male Long-Evans rats (350–400 g at the time of surgery) were housed individually in monitored conditions (21°C and 45% humidity) and maintained on a 12h light – 12h dark cycle. In order to avoid obesity, food was restricted to 13–16 g of rat chow per day, while water was available *ad libitum*. To habituate the rats to human manipulation, they were handled each workday. All experiments conformed to the approved protocols and regulations of the local ethics committee (Comité d’éthique en matière d’expérimentation animale Paris Centre et Sud n°59), the French Ministries of Agriculture, and Research.

### Surgery

The rats were deeply anesthetized with ketamine-xylazine (Imalgene 180 mg/kg and Rompun 10 mg/kg) and anesthesia was maintained with isoflurane (0.1-1.5% in oxygen). Analgesia was provided by subcutaneous injection of buprenorphine (Buprecaire, 0.025 mg/kg) and meloxicam (Metacam, 3 mg/kg). The animals were implanted with a custom built microdrive (144–252 channels) carrying 24, 32, or 42 independently movable hexatrodes (bundles of 6 twisted tungsten wires, 12 µm in diameter, gold-plated to ∼ 200 kΩ). The electrode tips were typically implanted 0.5 mm above the (bilateral) amygdala (– 2.4 mm to – 2.7 mm posterior to bregma and 4–5 mm lateral to bregma) and mPFC (0.3–0.6 mm lateral to bregma and 2.5–4.8 mm anterior to bregma). Miniature stainless steel screws were implanted above the cerebellum to serve as electrical reference and ground.

During recovery from surgery (minimum 7 days), the rats received antibiotic (Marbofloxacine, 2 mg/kg) and analgesic (Meloxicam, 3 mg/kg) treatments via subcutaneous injections and were provided with food and water *ad libitum*. The recording electrodes were then progressively lowered until they reached their targets and adjusted to optimize yield and stability.

## Data acquisition and processing

Brain activity was recorded using a 256-channel digital data acquisition system (KJE-1001, Amplipex, Szeged, Hungary). The signals were acquired with four 64-channel headstages (Amplipex HS2) and sampled wideband at 20,000 Hz. An inertial measurement unit (IMU, custom-made non-wireless version of the one described in (*69*)) sampled the 3D angular velocity and linear acceleration of the head at 300 Hz. To determine the instantaneous position of the animal, a red LED mounted on the headstage was imaged by overhead webcams at 30 Hz. Animal behavior was also recorded at 50 Hz by lateral video cameras (acA25000, Basler). Off-line spike sorting was performed using KiloSort (*70*) for prefrontal units, and KlustaKwik (K.D. Harris, http://klustakwik.sourceforge.net) for amygdalar units. The resulting clusters were visually inspected using Klusters (*71*) to reject noise and to merge erroneously split units. Neurophysiological and behavioral data were explored using NeuroScope (*71*). LFPs were derived from wideband signals by downsampling all channels to 1250 Hz.

## Scoring of behavioral and brain state

Automatic detection of immobility was performed by thresholding the angular speed calculated from gyroscopic data as described in (*69*). LFP data was visualized using Neuroscope (*71*) and slow-wave sleep (SWS) was detected as previously described (*72*).

## Histological identification of recording sites

At the end of the experiments, recording sites were marked with small electrolytic lesions (∼ 20 µA for 20 s, one lesion per bundle). After a delay of at least three days to permit glial scarring, rats were deeply anesthetized with a lethal dose of pentobarbital, and intracardially perfused with saline (0.9%) followed by paraformaldehyde (4%). Coronal slices (35 µm) were stained with cresyl-violet and imaged with conventional transmission light microscopy. Recording sites were reconstructed by comparing the images with the stereotaxic atlas of (*73*).

## Data analysis and statistics

Data were analyzed in Matlab (MathWorks, Natick, MA) using the Freely Moving Animal Toolbox (M. Zugaro and R. Todorova, http://fmatoolbox.sourceforge.net) and custom written programs. Detailed statistics are reported in Table S1.

## Identification of candidate cell assemblies

A standard unsupervised method based on principal and independent component analyses (PCA (*14*) and ICA (*25*), see (*74*)) detected the co-activation of simultaneously recorded neurons. Spike trains recorded during SWS were first binned into 15-ms bins and z-scored to generate a z-scored spike count matrix *Z*, where *Z_i_*_,*j*_ represents the activity of neuron *i* during time bin *j*. Principal components (PCs) were computed by eigen decomposition of the correlation matrix of *Z*. Principal components associated with eigenvalues exceeding the upper bound of the Marčenko-Pastur distribution were considered significant (*75*). We then carried out ICA (using the fastICA algorithm by H. Gävert, J. Hurri, J. Särelä, and A. Hyvärinen, http://research.ics.aalto.fi/ica/fastica) on the projection of *Z* onto the subspace spanned by significant PCs. Independent component (IC) weights were scaled to unit length and by convention the arbitrary signs of the weights were set so that the highest absolute weight was positive. Members of candidate cell assemblies were identified using Otsu’s method (*76*) to divide the absolute weights into two groups maximizing inter-class variance, and neurons in the group with greater absolute weights were classified as members. Goodness of separation was quantified using Otsu’s effectiveness metric, namely the ratio of the inter-class variance to the total variance. This procedure yielded a set of vectors *C_i_* representing the candidate cell assemblies.

In these vectors, coactive neurons correspond to weights of the same sign. In theory, two groups of anti-correlated neurons that inhibit each other could be detected as a vector with both positive and negative weight members (‘mixed-signs’ assembly). However, the reverse is not necessarily true: in our dataset, mixed-signs assemblies did not correspond to anti-correlated groups of coactive neurons, but to larger ensembles of members with poor separation quality compared to same-sign assemblies (Fig. S13). This suggested that mixed-signs assemblies resulted from limitations of the ICA method in identifying independent components from the PCs (*25*). We therefore discarded mixed-signs assemblies from further analyses.

Since the theoretical framework of this study precludes validation of candidate cell assemblies by their correlation to behavioral variables (behavioral correlates), we used a cross-validation method. The data were split in two balanced sets of random time intervals and candidat assemblies were independently detected in the two halves of the data. Assemblies across the two halves were significantly correlated in the vast majority of cases (99%; Fig. S15).

## Peer prediction

Population coupling of assembly members was verified by quantifying to what extent the spiking activity of one member could be predicted from the spiking activity of all other members (*12*). For cross-validation, spike trains were divided into two non-overlapping partitions. Using one partition (‘training set’), for each assembly member *i*, a generalized linear model (GLM) was trained to predict its activity *Z_i_* from the activity of all other members of the same assembly. To test performance, the GLM prediction error was computed on the remaining partition (‘test set’). This procedure was repeated exchanging the training and testing sets, resulting in two-fold cross-validation. The quality of the prediction was assessed by comparing the median prediction error *e* to the median error *e_shuffled_* obtained by shuffling 50 times the predictions relative to the observed activity *Z_i_*. The prediction gain *g* was defined as *g* = *e_shuffled_/e −* 1 (*77*).

## Assembly activations

To study downstream responses to assemblies, we computed an instantaneous assembly activation strength:

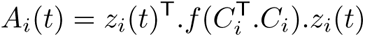

where *C_i_* contains the weights of the members of the *i^th^* assembly, and *z_i_*(*t*) is the activity of the assembly members at time *t* (computed using 15-ms windows and a 1-ms sliding window), and 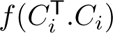 is a transformation of the outer product where the diagonal is set to 0, so that spiking in a single neuron does not contribute a high activation strength. Note that only the activity of assembly members were used in this computation to ensure that the activation strength reflects periods of coactivity of the assembly members rather than global fluctuations in the activity of cells with low weights (see Fig. S14). Assemblies were considered to be active when their activation strength exceeded a threshold of the 95th percentile of the values above baseline (the median, corresponding to empty bins). The midpoint of each threshold-exceeding activation was taken as assembly activation peak for further analyses.

## Downstream responses to candidate cell assemblies

For each candidate reader cell *i*, we computed the peri-event time histogram (PETH) of its spikes in the 2s interval (10 ms bins) centered on assembly activation peaks. PETHs with fewer than 30 spikes were discarded. To make computations tractable, candidate assembly–reader pairs were pre-selected for further analyses if the z-scored response exceeded 2 SDs in the 10–30 ms window following assembly activations (corresponding to the ∼20-ms conduction delay between these structures (*29*)). For each candidate assembly–reader pair, the response matrix was shuffled 200 times to determine pointwise and global confidence intervals (*13*). The pair was retained for further analysis if the following criteria were met: 1) the PETH was significant in at least one bin within the 10–30 ms window (crossing both the global and pointwise bands), and 2) the mode of the PETH was positive (the reader was activated after the assembly).

## Supralinearity of reader responses to assembly activations

To assess the supralinearity of reader responses to assembly activations, we first estimated the response that could be expected from a hypothetical reader responding independently to individual assembly members. To this end, we trained a generalized linear model (GLM) to predict the reader activity around assembly member spikes outside of assembly activations:

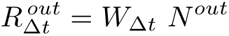

where *N^out^* is a (*m* + 1)-by-*n^out^* matrix containing the spike counts of each of the *m* assembly members (plus one constant term) in 15-ms bins around each of the *n^out^* assembly member spikes outside assembly activations, and 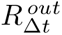 is a 1-by-*n^out^* vector containing the number of spikes of the reader neuron with a delay Δ*t* around each of the *n^out^* spikes; *W*_Δ*t*_ is a 1-by-(*m* + 1) vector containing the weights of the GLM fit for delay Δ*t* (Δ*t* varies between 1 s and 1 s) to produce the curves in Fig. 2. This linear model therefore captured the response of the reader at delay Δ*t* if the reader were responding to each individual assembly member independently. To estimate what the response of such a linear reader would be during assembly activations, we computed:

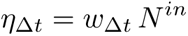

where *N^in^* is a (*m* + 1)-by-*n^in^* matrix containing the spike counts of each of the *m* assembly members (plus one constant term) in 15-ms bins around each of the *n^in^* assembly member spikes emitted during assembly activations, and *η*_Δ*t*_ is the activity predicted by the model for delay Δ*t*. Thus, the *collective* impact of the upstream assembly (beyond the sum of individual contributions) would be reflected in reader responses beyond *η*_Δ*t*_. We quantified this supralinearity by computing:

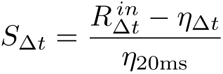

where 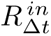 is a 1-by-*n^in^* vector containing the number of spikes of the reader neuron with a delay Δ*t* around each of the *n^in^* spikes, *η*_20ms_ is a normalisation factor corresponding to the estimated linear response *η*_Δ*t*_ at Δ*t* = 20 ms, and *S*_Δ*t*_ is the reader supralinearity at delay Δ*t*.

## Simulated readers

To estimate the supralinearity that would be expected from readers selective to collective activity vs unresponsive to collective activity, we repeated the above analyses on simulated data. We first simulated a ‘perfect’ reader that responded exclusively to the collective activity of the assembly: it only fired 20 ms after each partial activation recruiting at least half of the largest subset of co-active members (see *Pattern Completion* section below). We then simulated an ‘independent’ reader which fired 20 ms after every spike emitted by an assembly member, regardless of any collective activity.

## Time scales

The above analysis used a time scale of 15ms for cell assemblies (Fig. 2 and Fig. S6). We repeated this analysis for multiple time scales (Fig. S8). Candidate cell assemblies were detected as described above, but using time bins of 1 ms, 5 ms, 10 ms, 15 ms, 20 ms, 25 ms, 30 ms, 40 ms, 50 ms, 75 ms, and 100 ms. One critical issue with this analysis is that larger time bins may contain assemblies expressed at faster time scales: for instance, a 15-ms assembly also fits (and could thus also be detected) in time bins of e.g. 30ms or 100 ms — actually, in any time bin larger than 15 ms. To ensure that the analysis for a given time scale only used assemblies specifically expressed at that time scale, we excluded all epochs that contained activations of the same assembly at briefer time scales. The remaining activations were used to split member spikes between *n^in^* (assembly member spikes emitted during assembly activations) and *n^out^* (assembly member spikes emitted outside assembly activations). The two response curves (*R^in^* and *η*) were normalized conjointly: they were concatenated into a single vector for z-scoring.

## Selectivity to identity of assembly members

To test whether the reader was sensitive to the co-activation of multiple assembly members, rather than simply responding to the total spike output of assembly members, we compared the reader activity around co-activations of two different assembly members (‘AB’) to the reader activity around the repeated activation of a single assembly member (‘AA’).

For each assembly, we considered every possible permutation of two members. Each of these permutations was analyzed independently, and from herein, the two neurons in a given permutation are termed ‘A’ and ‘B’. To find ‘AB’ events, we performed a search in the inter-spike intervals of ‘A’ and ‘B’, retaining pairs of spikes emitted by the two neurons within the assembly time scale (15ms). To find a matching set of ‘AA’ events, we performed an equivalent search of moments when neuron ‘A’ emitted two consecutive spikes within the same time scale (15 ms). Permutations in which we found less than 20 ‘AB’ events or less than 20 ‘AA’ events were discarded from further analyses. We computed a peri-event time histogram (PETH) of the firing rate of the reader neuron around ‘AB’ and ‘AA’ events, using the midpoint of the two spikes (‘AB’ or ‘AA’) as a reference. The two PETHs were normalized conjointly: they were concatenated into a single vector for z-scoring.

The above analysis used a time scale of 15-ms (Fig. S7). We repeated the analysis for multiple time scales (Fig. 2, Fig. S9). Candidate cell assemblies were detected as described above, but using bins of 1 ms, 5 ms, 10 ms, 15 ms, 20 ms, 25 ms, 30 ms, 40 ms, 50 ms, 75 ms, and 100 ms. For each time scale, we detected candidate readers using the procedure outlined above. We further subdivided reader responses according to the delay between the two spikes, within a precision of 5 ms. For example, to compute the reader response to ‘AA’ events with a delay of 45 ms, we retained ‘AA’ events for which the two consecutive spikes were within 42.5–47.5 ms of each other (without any ‘A’ or ‘B’ intervening spikes during this interval).

## Pattern Completion

To quantify pattern completion, we determined the average reader responses to activation of all possible combinations of assembly members. For example, for assembly ‘ABCD’, we measured reader responses to the (complete) 4-member assembly activations ‘ABCD’, to each of the 3-member (incomplete) activations ‘ABC’, ‘ABD’, ‘ACD’, ‘BCD’, to each of the 2-member (incomplete) activations ‘AB’, ‘AC’, ‘AD’, ‘BC’, ‘BD’, ‘CD’, and to each of the single-member (incomplete) activations ‘A’, ‘B’, ‘C’, ‘D’, relative to the baseline reader firing rate in all sleep periods. For each assembly–reader pair, we fit the resulting responses with a sigmoid curve:

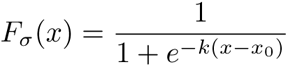

where *x* is the proportion of active assembly members, and *x*_0_ and *k* are the model parameters corresponding to the midpoint and the steepness of the curve, respectively. To estimate the goodness-of-fit, we computed

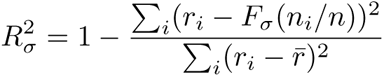

where *r_i_* is the reader response to combination *i*, and *n_i_/n* is the proportion of active members. We likewise estimated the goodness-of-fit of a proportional response *F_α_*(*x*) = *r_complete_ x*, where *r_complete_* is the reader response to activations of the complete assembly (or of the largest subset of co-active members):

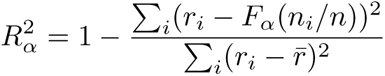

Finally, we quantified the boost in observed response relative to the proportional response as the gain *r – F_α_*. We split the data in tertiles according to *x* such that *x*_1_ ∊ (0, 1*/*3], *x*_2_ ∊ (1*/*3, 2*/*3], and *x*_3_ ∊ (2*/*3, 1] and tested each tertile for significant pattern completion.

### Pattern Separation

To assess how readers discriminated between different assemblies, we first determined the response of each neuron *j* following activations of each recorded assembly *i*, and computed the Hoyer coefficient of sparsity:

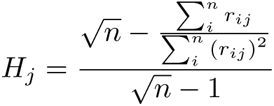

where *n* is the number of assemblies recorded simultaneously with neuron *j*, and *r_ij_* is the response of neuron *j* to assembly *i*. As neurons with lower baseline firing rates tend to have larger Hoyer coefficients of sparsity, to compare across neurons we measured sparsity relative to surrogate data, where assembly identities were shuffled across all pooled assembly activations (i.e., an activation of assembly *a* was randomly assigned to assembly *b*). For each reader, we repeated this procedure 1000 times and computed the mean Hoyer coefficient of the shuffled data 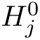. The sparsity increase relative to the shuffled data was defined as:

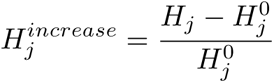

To determine whether reader responses were particularly sparse, we compared sparsity increases between readers and non-readers (neurons for which a paired assembly could not be detected) using the Wilcoxon rank sum test.

To test whether readers could discriminate between similar patterns, for each reader–assembly pair we sought a second assembly with multiple overlapping members (at least 25% of each assembly and *>* 2 members, e.g. ‘ABCD’ and ‘ABE’; varying the number of overlapping members did not change our results: see Supplementary Figure 11), and defined the discrimination index between the two assemblies as:

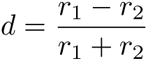

where *r*_1_ is the reader response to its paired assembly, and *r*_2_ is its response to the overlapping assembly. To test for significant discrimination, we computed discrimination indices for surrogate data, where the activations of the two assemblies were pooled and the assembly identities were shuffled. This was repeated 1000 times, and when the discrimination index of a reader exceeded those of 95% of the shuffles, the reader was considered to perform pattern separation.

## Behavioral testing

Behavioral testing only relates to results shown in Fig. 5 as all other analyses were performed using data from pre-training sleep sessions, i.e. preceding any exposure to the protocol described here.

The animals were tested in a slightly extended version of the standard fear conditioning and extinction paradigm, initially intended to discriminate between cued and context fear learning. However, only context extinction yielded useful data for the current study and is reported here. Briefly, fear conditioning took place in one chamber (context), where foot shocks were associated with auditory stimuli (conditioned stimuli, CS). Extinction took place either in the same chamber without the CS (contextual extinction), or in a different chamber with the CS (cued extinction). Daily recording sessions consisted of two 37-min exposure sessions (one per chamber), preceded, separated, and followed by sleep sessions of 2-3 hours. Only sleep periods before and after exposure to the conditioning chamber were analyzed here, amounting to two pre-training control sessions, two fear conditioning sessions, and two extinction sessions per animal.

The conditioning chamber was cubic (side length, 40 cm) with gray plexiglass walls lined with ribbed black rubber sheets and a floor composed of nineteen stainless steel rods (0.48 cm diameter, 1.6 cm spacing) connected to a scrambled shock generator (ENV-414S, Med Associates, USA). It was mildly scented daily with mint-perfumed cleaning solution (Simple Green, Sunshine Makers). A custom-made electronic system presented the animals with two auditory CS (80 dB, 20 s long, each composed of 1 Hz, 250 ms long pips of either white noise, CS+ paired to shocks, or 8 kHz pure tones, CS-unpaired). These auditory stimuli (8 CS+ and 8 CS-) were presented starting at *t* = 3 min, separated by random-duration inter-trial intervals (120–240 s). Foot shocks consisted in shocks scrambled across floor rods (1 s, 0.6 mA, co-terminating with CS+ presentations; CS+ and CS- were presented in pseudorandom order allowing no more than 2 consecutive presentations of the same-type CS). Sleep was recorded in a cloth-lined plastic flowerpot (30 cm upper diameter, 20 cm lower diameter, 40 cm high).

## Data availability

The datasets generated during the current study are available in the [NAME] repository [LINK WILL BE PROVIDED UPON ACCEPTATION].

---

**Fig. S1.**
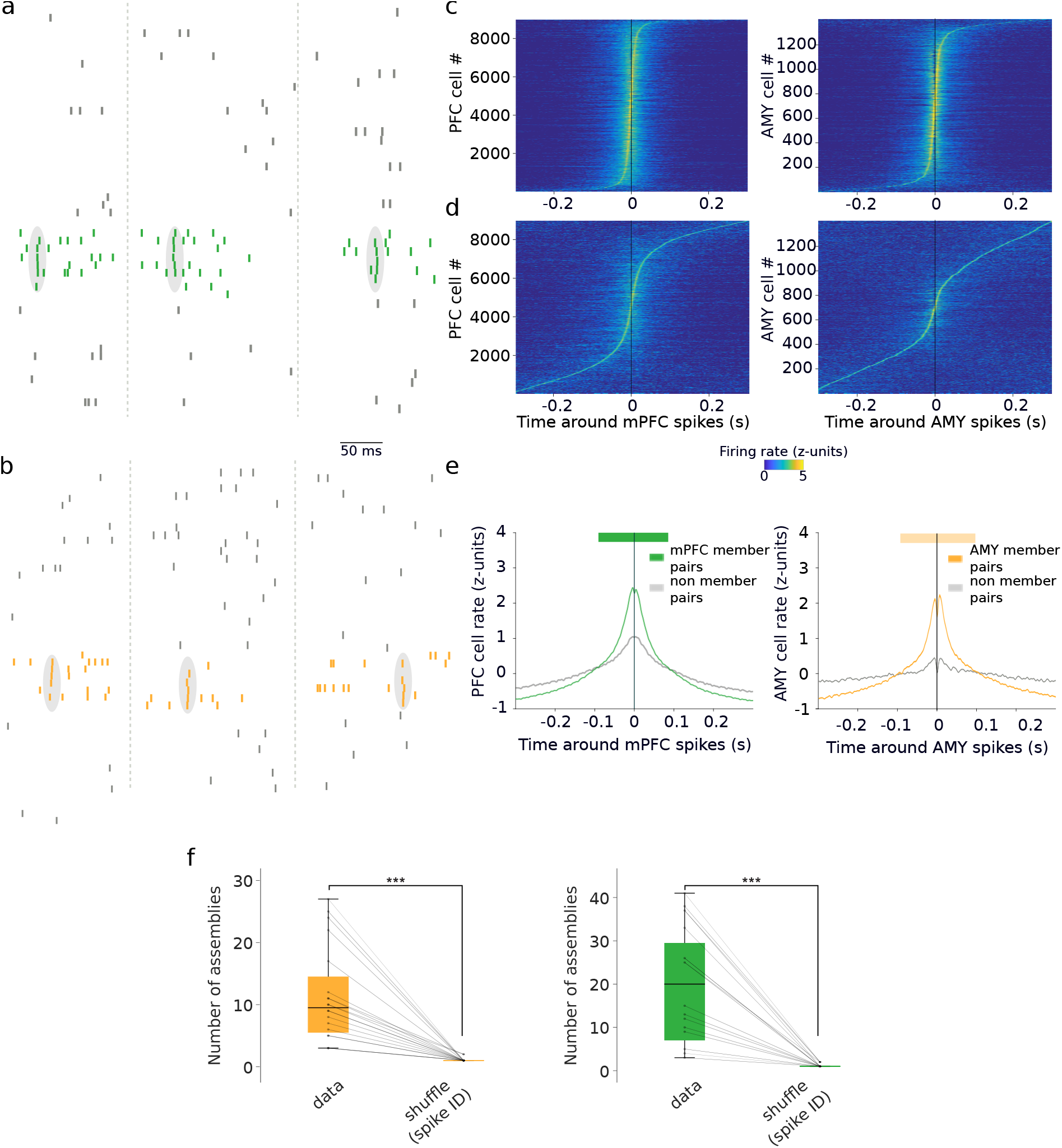
Candidate cell assemblies in the cortico-amygdalar circuit. **a**, Spike trains of a subset of 35 simultaneously recorded units in prefrontal cortex during sleep (rasters: action potentials; gray ellipses surrounding colored ticks: co-activation events). **b**, Spike trains of a subset of 30 simultaneously recorded units in the amygdala during sleep (rasters: action potentials; gray ellipses surrounding colored ticks: co-activation events). **c**, Z-scored cross-correlations between members of the same prefrontal (left) and amygdalar (right) assemblies, ordered by mode. **d**, Same as in (**c**) for control pairs, illustrating that fewer pairs have modes at brief delays. **e**, Averages of (**c**) (colored curves) and (**d**) (gray curves). Members of the same assemblies had significantly higher synchrony at short delays than control pairs (thick horizontal colored bars: *p <* 0.05, Monte-Carlo bootstraps). **f**, Number of candidate assemblies in the amygdala (left) and mPFC (right) in actual vs. shuffled data preserving global rate fluctuations.

**Fig. S2.**
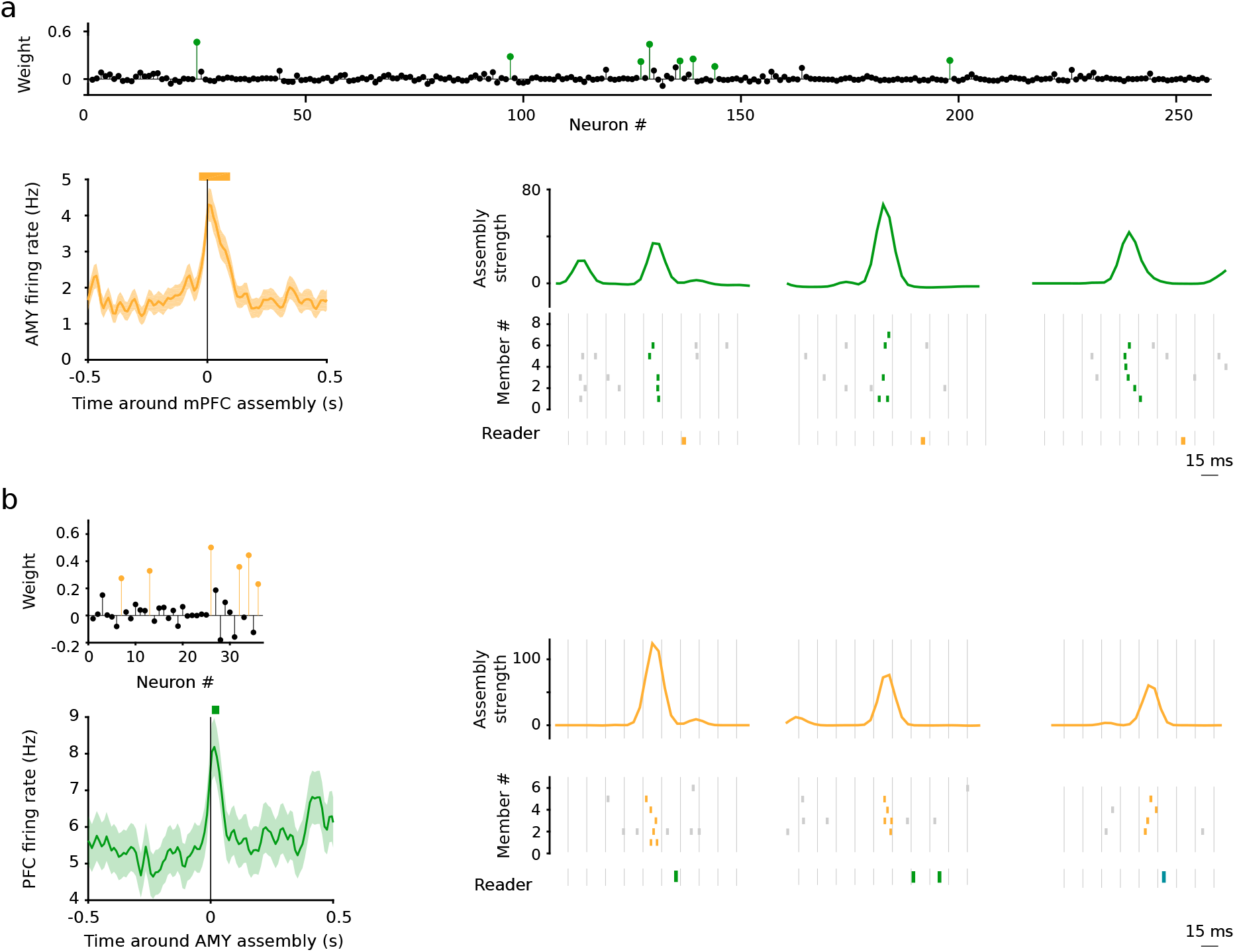
Example assembly–reader pairs. **a**, Activations of a prefrontal assembly closely followed (10– 30 ms) by significant responses of an amygdalar neuron. Top: cell assembly weights (colored circles: assembly members, black circles: non-members). Bottom left: firing rate of an amygdalar neuron centered on all prefrontal assembly activations (mean ± s.e.m.). Thick orange horizontal bar indicates significant responses (*p <* 0.05: Monte-Carlo bootstrap test; see Methods). Bottom right: example assembly activations (green curves: activation strength) followed by downstream spiking (rasters: prefrontal spikes within (green) or outside (gray) epochs of assembly activation; orange rasters: amygdalar spikes). Reader responses occurred ~20 ms after assembly activations. **b**, Same as (**a**) for an amygdalar assembly and a downstream prefrontal neuron.

**Fig. S3.**
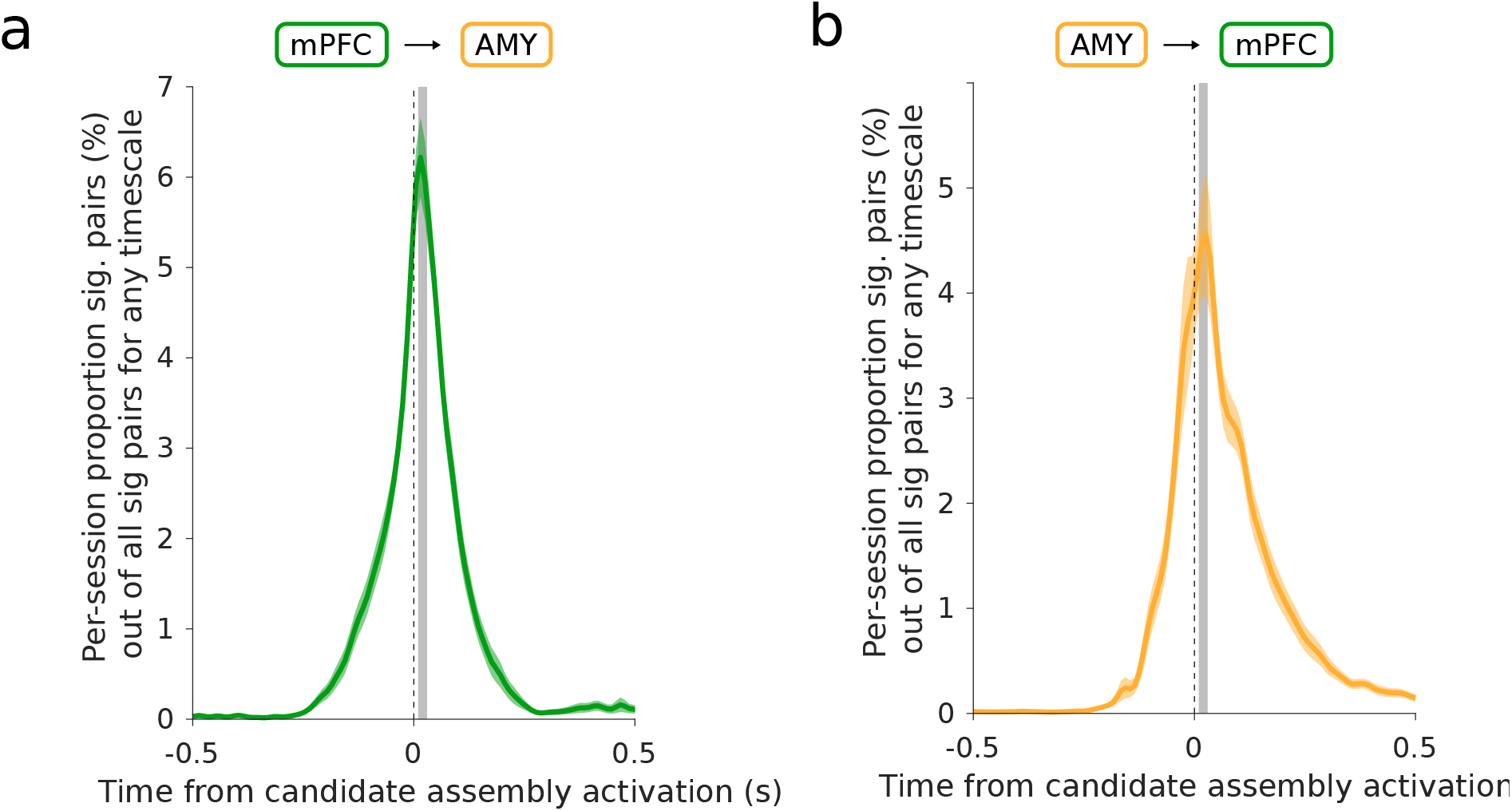
Optimal delay for detecting assembly–reader pairs. **a**, Percentage of significant pairs of candidate prefrontal assemblies and amygdalar readers as a function of the assembly–reader delay. Note that the number of significant pairs peaked for amygdala readers responding ∼ 20 ms after candidate prefrontal assembly activation. **b**, Same as (**a**) for candidate amygdalar assemblies and prefrontal readers.

**Fig. S4.**
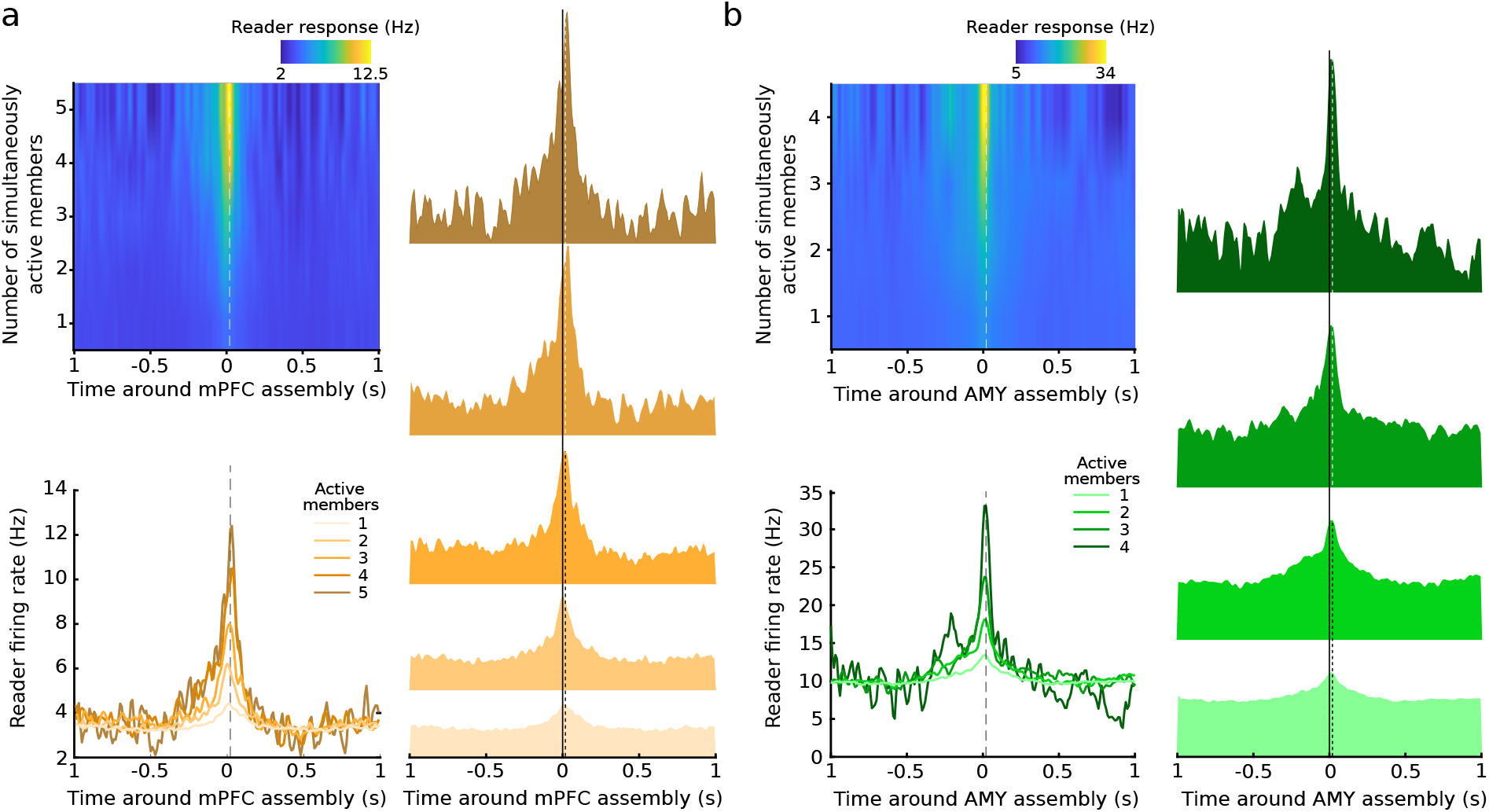
Assembly members exert a synergistic influence on their targets: reader response rate increases with the number of co-active assembly members. **a**, Example amygdalar response to increasing numbers of simultaneously active prefrontal assembly members. Top left: Reader firing rate centered on assembly activation. Right: Reader firing rate for different numbers of co-active members. Bottom left: Superimposed response curves. **b**, Same as (**a**) for example prefrontal assembly and amygdalar reader.

**Fig. S5.**
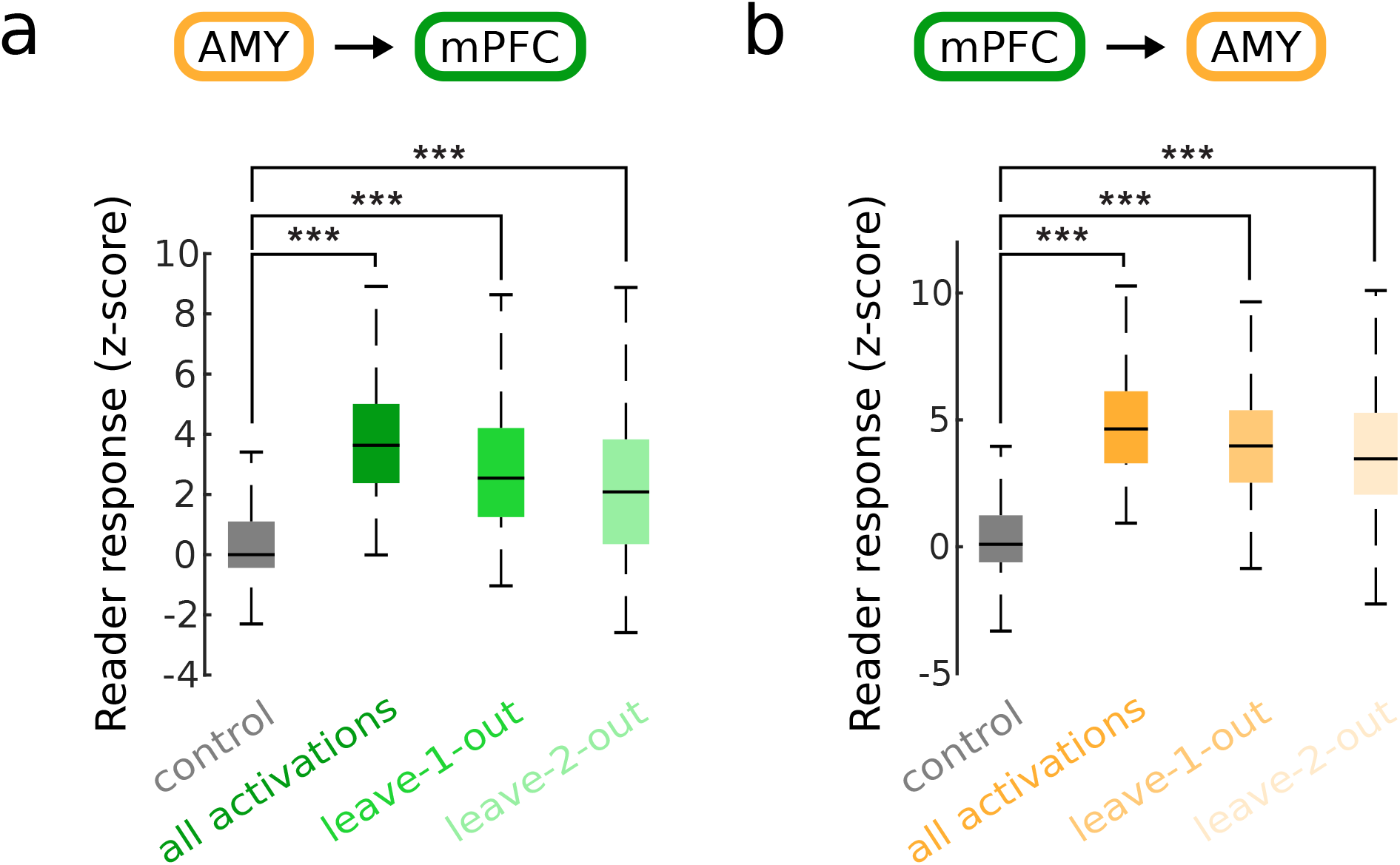
Assembly members exert a synergistic influence on their targets: responses are not driven by single effective members. **a**, Average response of prefrontal readers to amygdalar assembly activations when the most effective members (i.e. the members whose spikes outside assembly activation epochs were followed by the largest response by the reader neuron at 10–30 ms) of upstream assemblies were not recruited (leave 1-out, leave 2-out). **b**, Same as (**a**) for amygdalar reader responses to member spikes of prefrontal assemblies.

**Fig. S6.**
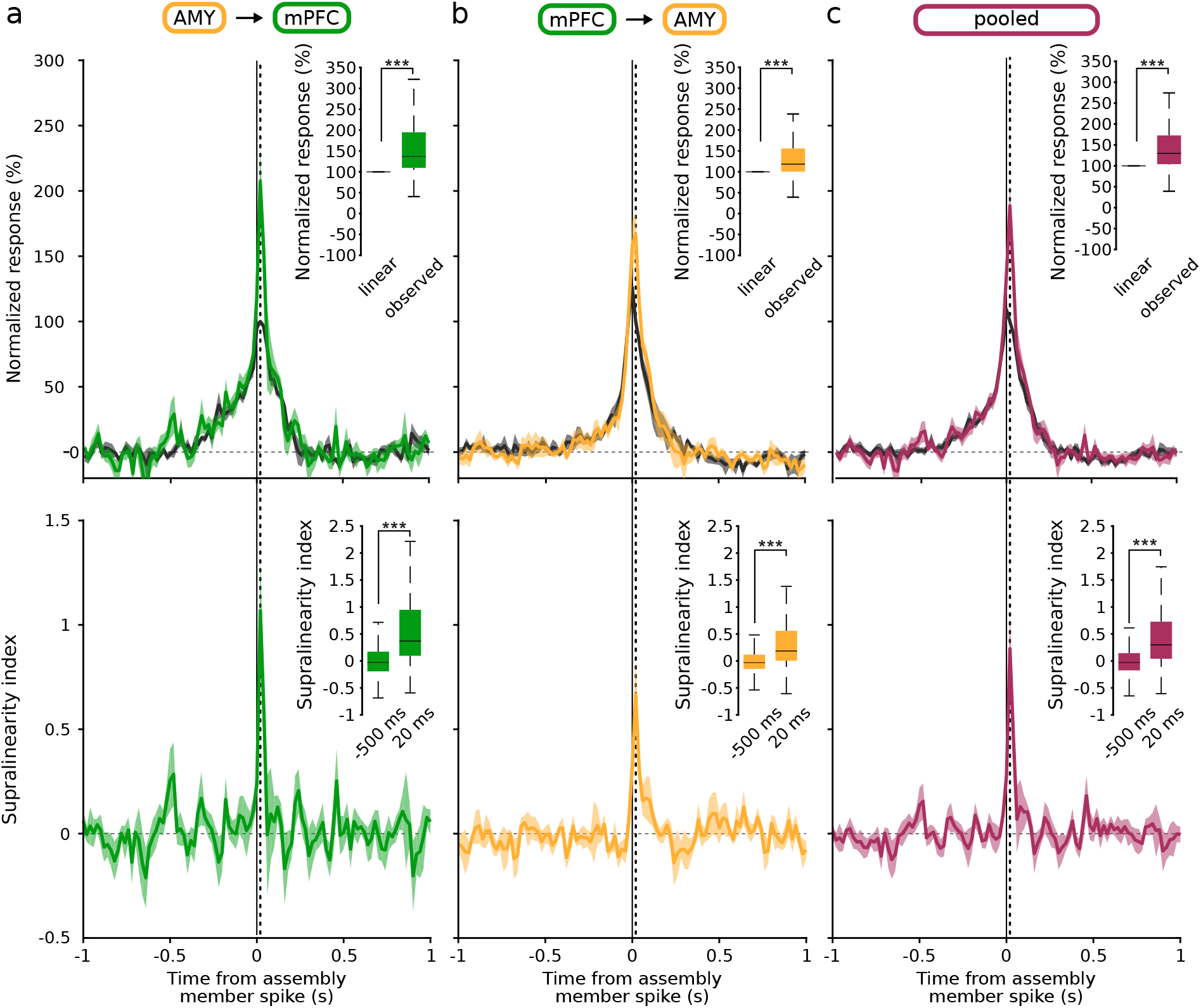
Supralinearity of reader responses. **a**, Top: Observed responses (colored curve: mean ± s.e.m.) of prefrontal readers compared to the estimated response of a linear reader (gray curve: mean ± s.e.m.). Inset: The observed response was greater than the linear estimate at 20 ms (\*\*\**p <* 0.001, Wilcoxon signed-rank test). Bottom: Supralinearity index of prefrontal reader responses. Dashed line: peak of reader responses to assembly activations at 20 ms. Inset: Supralinearity at 20 ms vs baseline (\*\*\**p <* 0.001, Wilcoxon signed-rank test). **b**, Same as (**a**) for amygdalar reader responses to spikes of members of prefrontal assemblies. **c**, Same as (**a**) for pooled responses of both amygdalar and prefrontal readers.

**Fig. S7.**
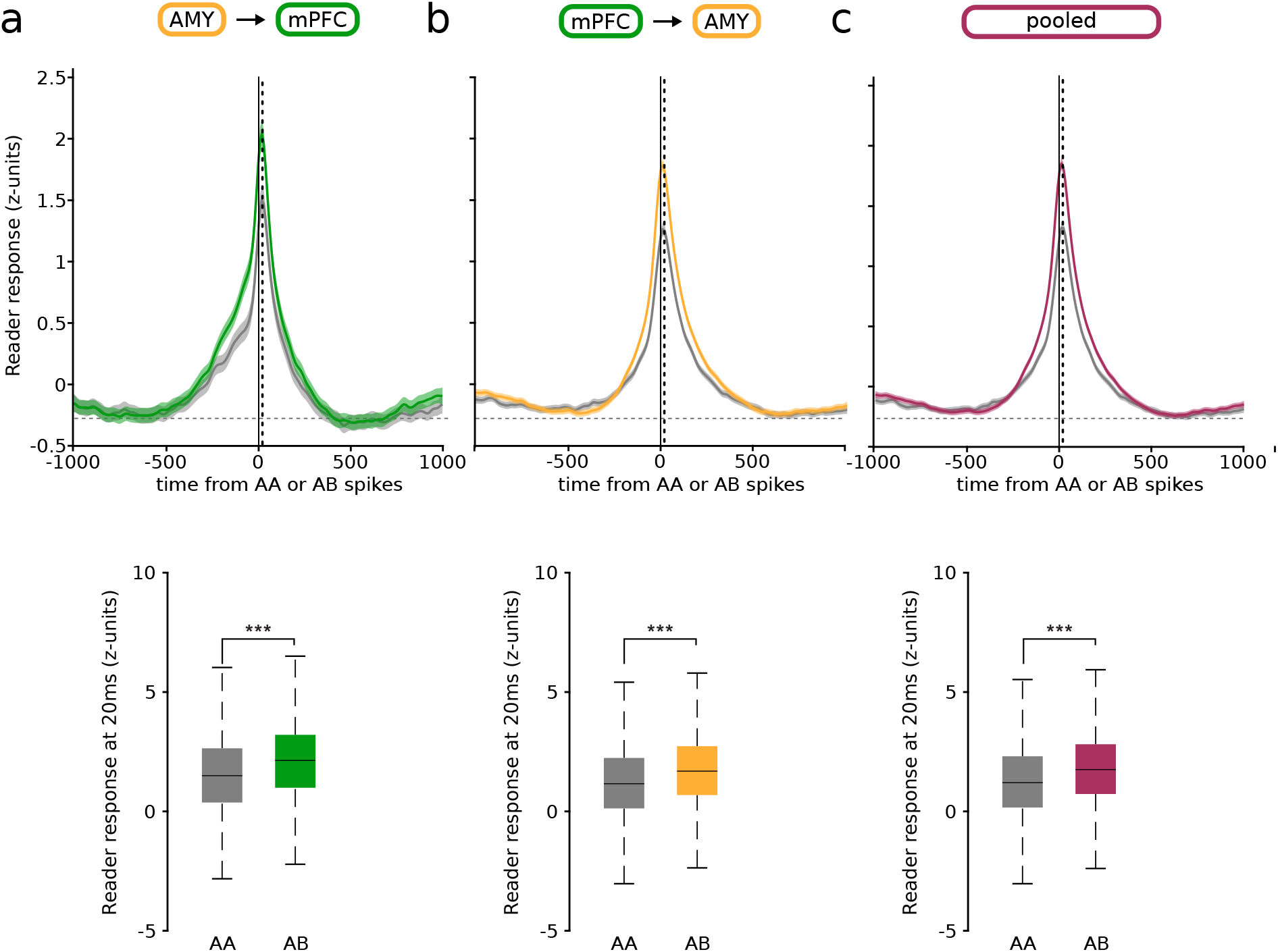
The identity of participating members matters beyond their compound activity. **a**, Response of prefrontal readers to amygdalar assembly members. Top: z-scored responses of reader neurons to two spikes emitted by different assembly members (AB, colored curve), compared to the control responses to two spikes emitted by the same assembly member (AA, gray curve) (mean ± sem). Bottom: Z-scored reader responses at 20 ms (\*\*\**p <* 0.001, Wilcoxon signed-rank test). **b**, Same as (**a**) for amygdalar readers and PFC assembly members. **c**, Same as (**a**) for pooled responses of both amygdalar and prefrontal readers.

**Fig. S8.**
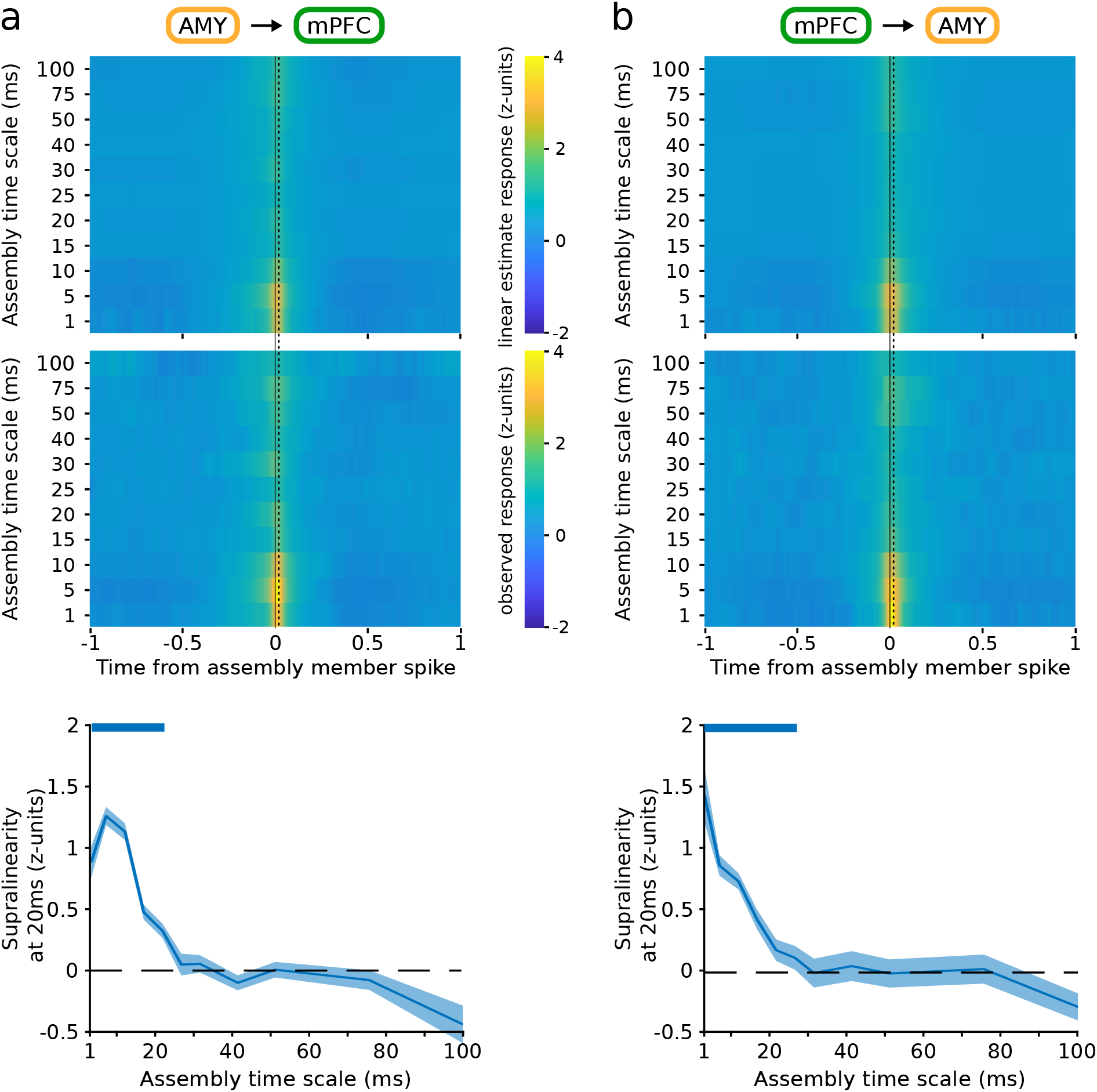
Time scale of reader response supralinearity. **a**, Response of prefrontal readers to activations of amygdalar assemblies at varying time scales. Top and center: mean z-scored responses of a linear model vs the observed reader response, as a function of the time scale of the assembly. Bottom: difference between the two (observed response linear estimate), for varying time scales. Thick colored horizontal bars indicate significant differences (*p <* 0.05, Monte-Carlo bootstrap test). **b**, Same as (**a**) for amygdalar reader responses to member spikes of prefrontal assemblies. Note that in both cases, supralinearity is significantly greater than 0 for time scales up to 20–25 ms.

**Fig. S9.**
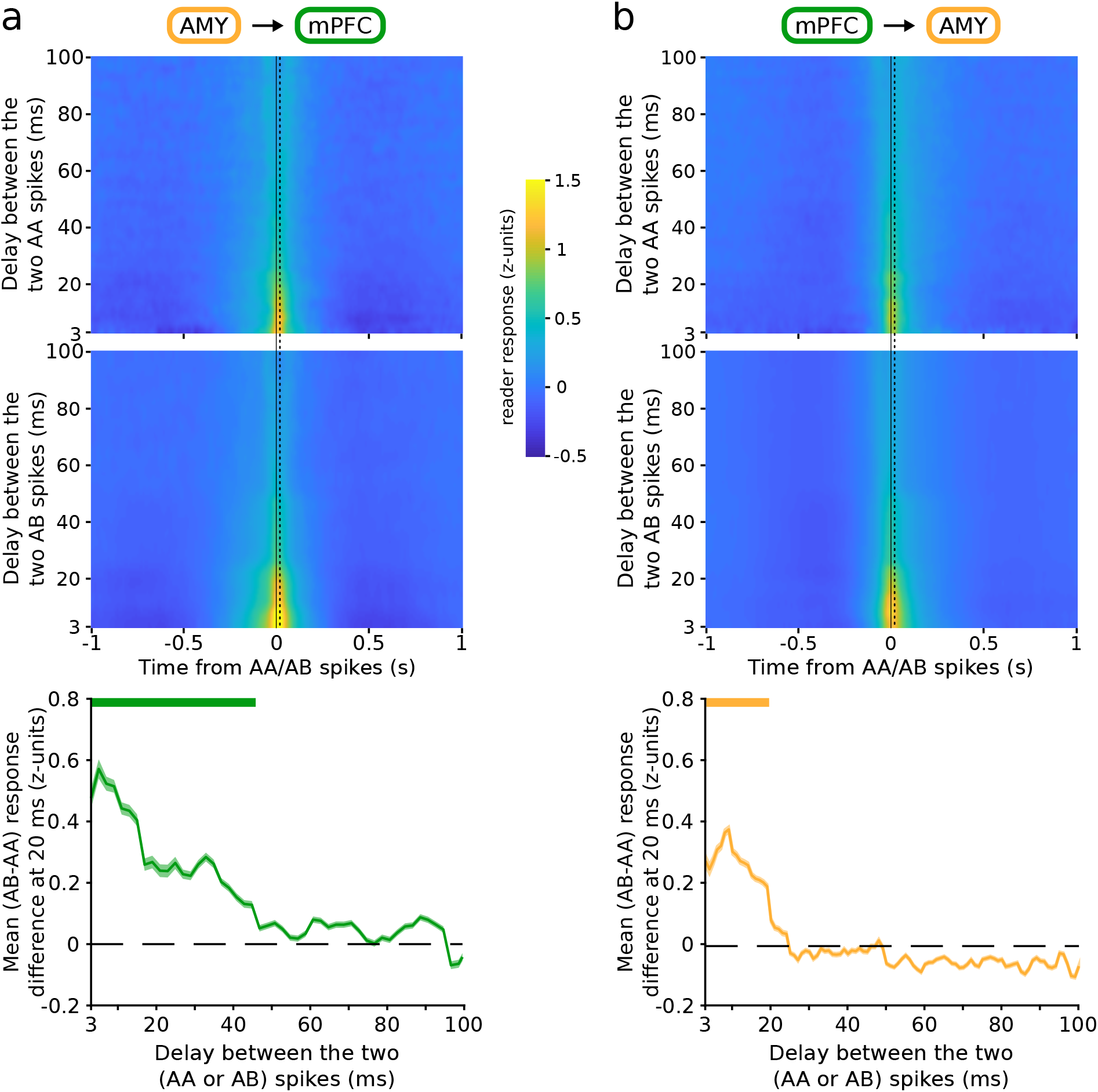
Time scale of reader sensitivity to assembly member identity. **a**, Response of prefrontal readers to two spikes emitted by different assembly members (AB), compared to the control responses to two spikes emitted by the same assembly member (AA), at varying time scales. Top and center: mean z-scored responses of reader neurons to spikes emitted by the same (AA, top) vs different (AB, center) members of an upstream assembly, as a function of the temporal delay between the two spikes. Bottom: difference between the two (AB AA), for varying temporal delays. Thick colored horizontal bars indicate significant difference (*p <* 0.05, Monte-Carlo bootstrap test). **b**, Same as (**a**) for amygdalar readers.

**Fig. S10.**
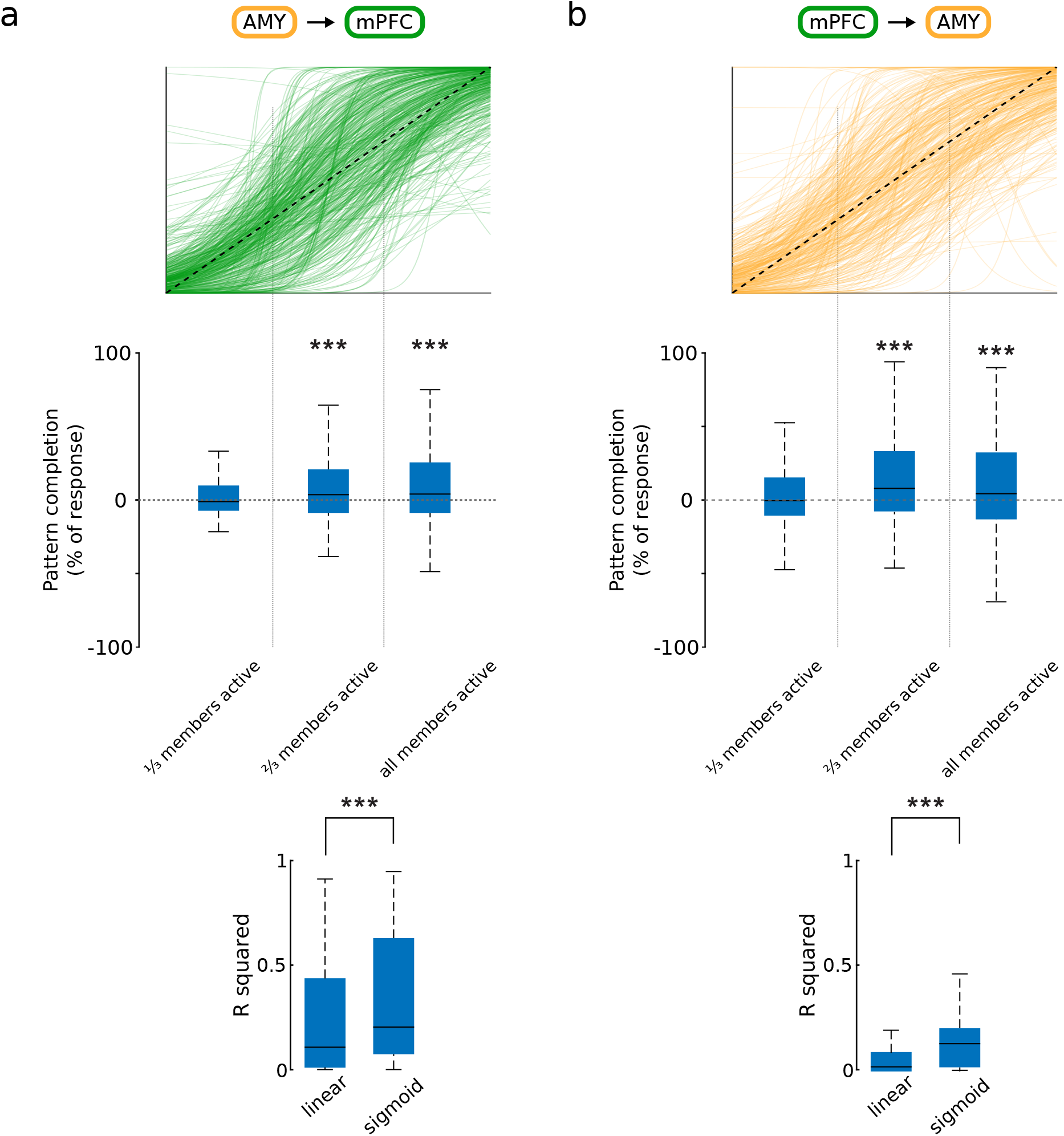
The assembly–reader mechanism can implement pattern completion. **a**, Prefrontal reader responses to amygdalar assemblies. Top: Superimposed best-fit sigmoid curves of all assembly–reader pairs. Center: boost in reader response (relative to a proportional response) for all assembly–reader pairs as a function of the proportion of active assembly members. The gain was significant for the second and third quantiles (\*\*\**p <* 0.001, Wilcoxon signed-rank test), but not for the first quantile (\**p <* 0.05, Wilcoxon signed-rank test). Bottom: The data were better fit with sigmoidal than linear models (\*\*\**p <* 0.001, Wilcoxon signed-rank test). **b**, Same as (**a**) for amygdalar reader responses to prefrontal assemblies.

**Fig. S11.**
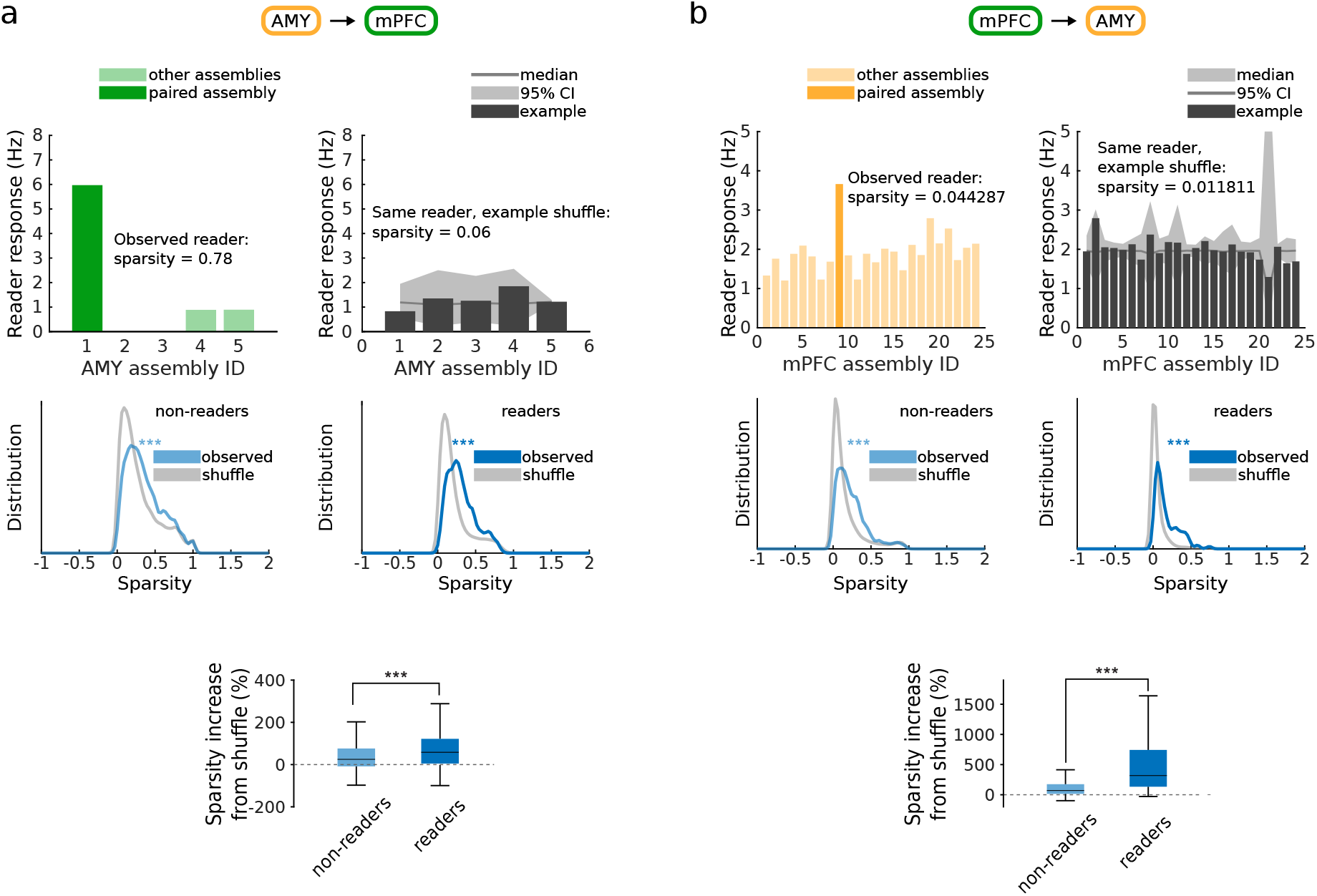
The assembly–reader mechanism can implement pattern separation: reader responses are selective for specific assemblies. **a**, Sparsity of prefrontal reader responses to amygdalar assemblies. Top left: Responses of an example prefrontal neuron to each amygdalar cell assembly in the recording session. Responses are selective for the paired assembly (dark green), compared to other assemblies (light green). Top right: Control responses of the same prefrontal neuron to surrogate assembly activations (shuffled assembly identities) are not selective. Center: distribution of sparsity for non-reader (left) and reader (right) neurons, compared to control sparsity computed from shuffled data (gray). Note that the observed responses are sparser than the shuffled control (\*\*\**p <* 0.001, Wilcoxon signed rank test). Bottom: Sparsity increase from shuffle, for reader vs non-reader neurons (\*\*\**p <* 0.001, Wilcoxon rank sum test). **b**, Same as (**a**) for amygdalar reader responses to prefrontal assemblies.

**Fig. S12.**
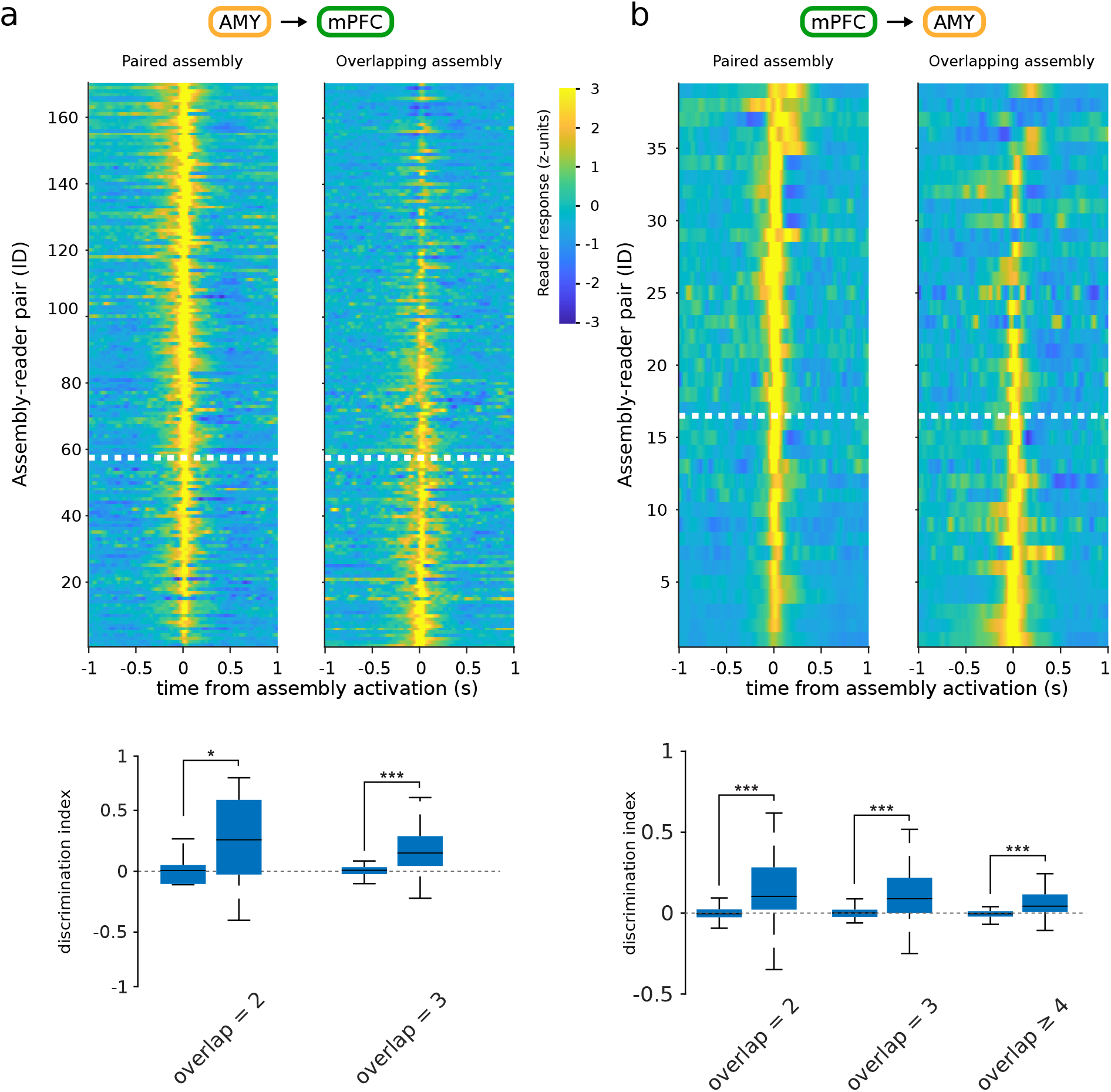
The assembly–reader mechanism can implement pattern separation: readers can discriminate between overlapping assemblies. **a**, Pattern separation in prefrontal reader responses to amygdalar assemblies. Top: prefrontal reader responses to activation of a paired assembly (left) vs a different but overlapping (≥ 25%) assembly, sorted by discrimination index. Responses above the white dotted line manifested significant pattern separation (greater discrimination indices than shuffled data, *p <* 0.05, Wilcoxon rank sum test). Bottom: Discrimination indices were greater for observed than shuffled data (\*\*\**p <* 0.001, Wilcoxon rank sum test). **b**, Same as (**a**) for amygdalar reader responses to prefrontal assemblies.

**Fig. S13.**
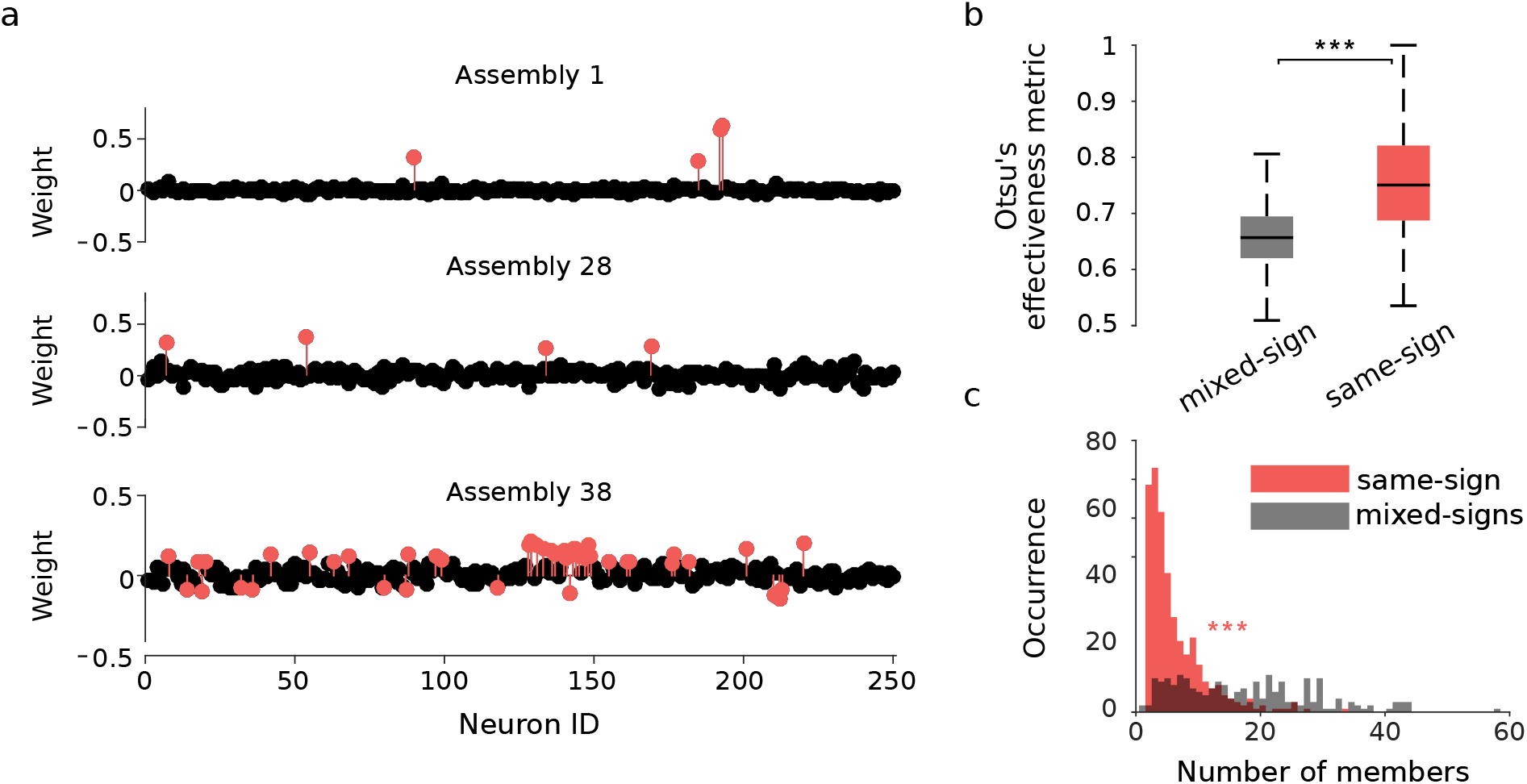
Selection of candidate cell assemblies with same-sign component weights. **a**, Cell assembly weights of three representative prefrontal assemblies (colored circles: assembly members, black circles: non-members), corresponding to eigenvalues 1, 28 and 38. Whereas all members of assemblies 1 and 28 were of the same (positive) sign, assembly 38 included members with both positive and negative weights (‘mixed-sign assembly’). **b**, The separation between members and non-members was significantly better in same-sign assemblies than mixed-sign assemblies (\*\*\**p <* 0.001, Wilcoxon rank sum test). **c**, Mixed-sign assemblies had significantly more members than same-sign assemblies (\*\*\**p <* 0.001, Wilcoxon rank sum test).

**Fig. S14.**
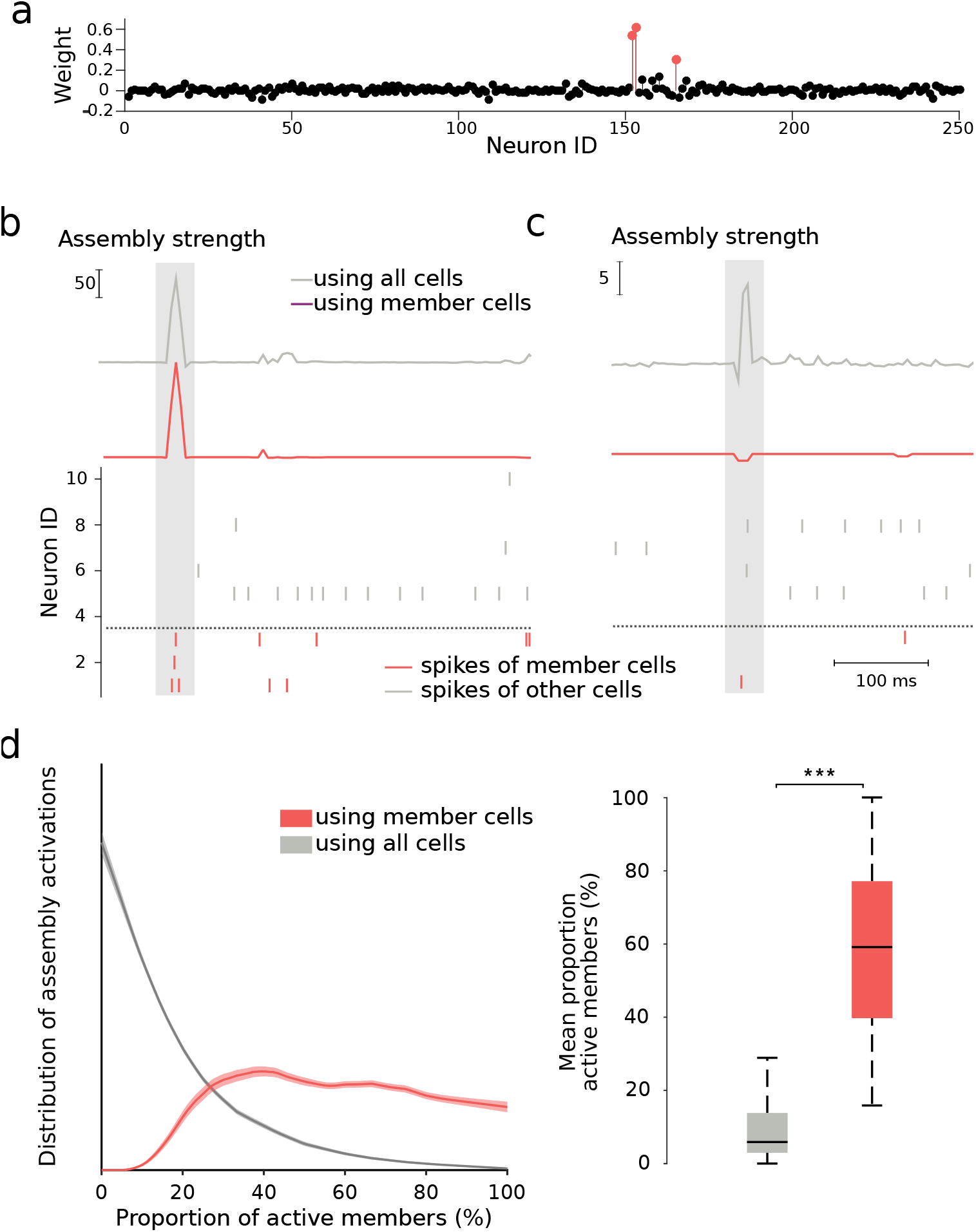
Computation of assembly activation strength. **a**, An example assembly recorded in the prefrontal cortex. Red dots: assembly members. **b**, Example activation of assembly shown in (**a**). Top: Assembly activation strength computed using either the activity of all cells (gray curve) or the activity of member cells only (red curve). Bottom: Raster plot of the activity of a representative subset of neurons, ordered by absolute weight (vertical ticks: action potentials; red ticks: member cells; gray ticks: non-member cells; shaded rectangle: putative assembly activation). All three members were active, resulting in a high activation strength in both curves. **c**, Same as (**b**) but for an instance in which only a single assembly member was active, at the same time as two non-members. The corresponding peak in the gray curve would result in incorrect detection of an activation of the assembly. This spurious peak is absent from the red curve, where activity strength is computed using only assembly members. **d**, Left: Proportion of assembly members co-active around peaks in the assembly activation strength computed using the activity of member cells only (red) or using the activity of all cells (gray). Right: using the activity of member cells results in detection of assembly activation events with greater proportions of co-active members (\*\*\**p <* 0.001, Wilcoxon signed rank test)).

**Fig. S15.**
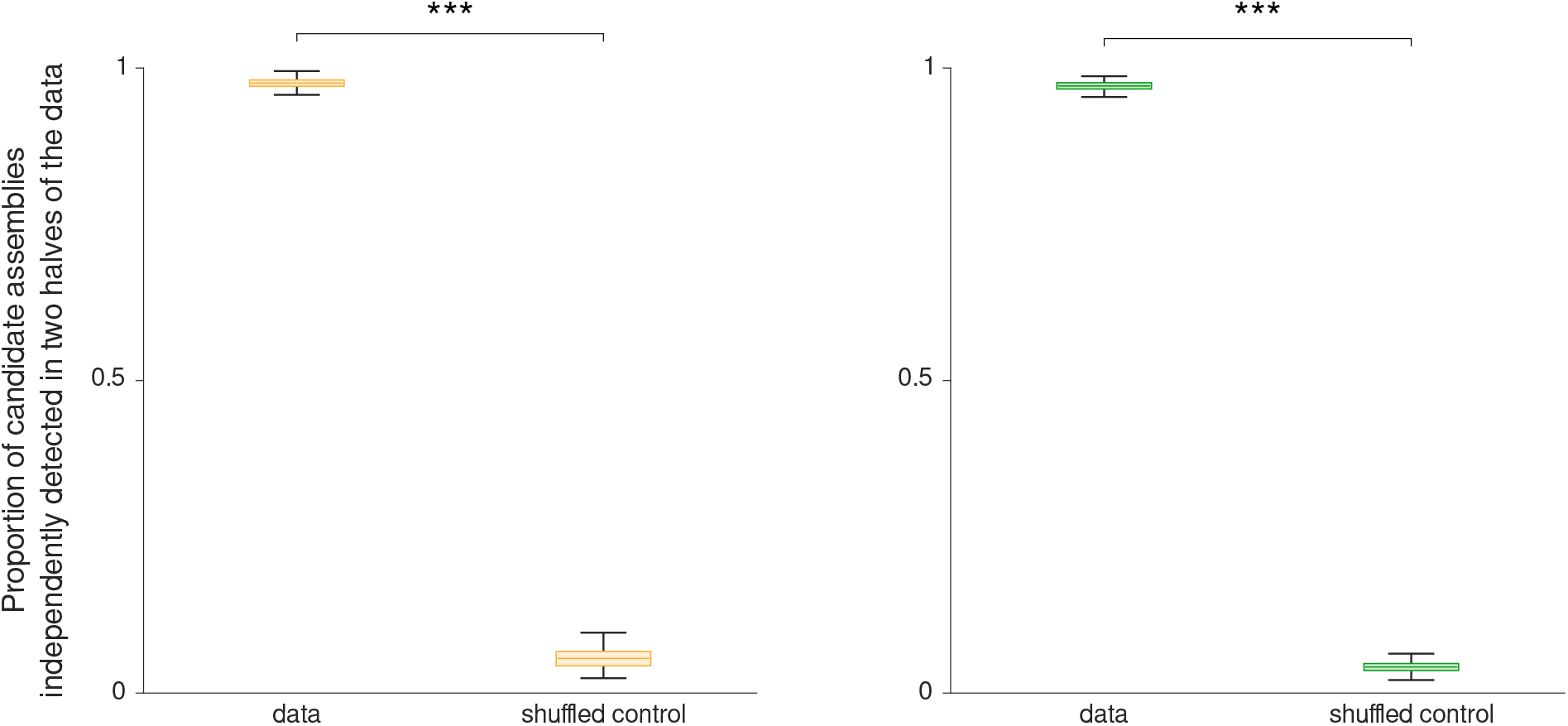
Cross-validation of detection of candidate cell assemblies. **a**, Proportion of candidate amygdalar assemblies that were independently detected in each half of the recorded data (\*\*\**p <* 0.001, Wilcoxon signed rank test). **b**, Same as (**a**) for candidate prefrontal assemblies.

**Table S1:** Detailed Statistics.

| Figure | Test | Description | P-value | N | Effect size |
| --- | --- | --- | --- | --- | --- |
| 1.b (left) | Wilcoxon rank sum test | GLM gain (%) vs shuffled data (PFC) | $p = 0$ | 2061 | ∞ |
| 1.b (right) | Wilcoxon rank sum test | GLM gain (%) vs shuffled data (AMY) | $p = 1.767 \cdot 10^{-202}$ | 676 | 30,3 |
| 1.f (top) | Wilcoxon signed-rank test | % of assembly-reader pairs vs chance | $p = 6.1 \cdot 10^{-5}$ | 20 | 4 |
| 1.f (bottom) | Wilcoxon signed-rank test | % of assembly-reader pairs vs chance | $p = 1.2 \cdot 10^{-4}$ | 20 | 3,84 |
| 2.a | Wilcoxon rank sum test | reader neuron responses (all activations) vs control | $p = 0$ | 849 | ∞ |
| 2.a | Wilcoxon rank sum test | reader neuron responses (leave-1-out) vs control | $p = 1.1887 \cdot 10^{-320}$ | 849 | 38,2827 |
| 2.a | Wilcoxon rank sum test | reader neuron responses (leave-1-out) vs control | $p = 2.0842 \cdot 10^{-196}$ | 849 | 29,8981 |
| 2.b | Wilcoxon signed-rank test, one-tailed | supralinearity index (observed data) vs baseline | $p = 1.336 \cdot 10^{-98}$ | 1173 | $z = 21.043$ |
| 2.b | Wilcoxon signed-rank test, one-tailed | supralinearity index (perfect reader) vs baseline | $p = 8.416 \cdot 10^{-169}$ | 1173 | $z = 27.668$ |
| 2.b | Wilcoxon signed-rank test, one-tailed | supralinearity index (independent reader) vs baseline | $p = 0.7916$ | 1173 | $z = -0.812$ |
| 2.c | Wilcoxon signed-rank test, one-tailed | response to AA events vs AB events (5 ms delay) | $p = 1.840 \cdot 10^{-91}$ | 4443 | $z = 20.248$ |
| 2.c | Wilcoxon signed-rank test, one-tailed | response to AA events vs AB events (10 ms delay) | $p = 3.088 \cdot 10^{-87}$ | 6520 | $z = 19.763$ |
| 2.c | Wilcoxon signed-rank test, one-tailed | response to AA events vs AB events (15 ms delay) | $p = 3.072 \cdot 10^{-121}$ | 10288 | $z = 23.384$ |
| 2.c | Wilcoxon signed-rank test, one-tailed | response to AA events vs AB events (20 ms delay) | $p = 2.490 \cdot 10^{-77}$ | 8487 | $z = 18.576$ |
| 2.c | Wilcoxon signed-rank test, one-tailed | response to AA events vs AB events (25 ms delay) | $p = 0.1315$ | 12056 | $z = 1.119$ |
| 2.c | Wilcoxon signed-rank test, one-tailed | response to AA events vs AB events (50 ms delay) | $p = 1$ | 10383 | $z = -9.900$ |
| 2.c | Wilcoxon signed-rank test, one-tailed | response to AA events vs AB events (100 ms delay) | $p = 0.9996$ | 8166 | $z = -3.415$ |
| line 102 | chi-square test | amygdalar readers vs non-readers likeliness to participate in candidate cell assemblies targeting prefrontal readers | $p = 1.2 \cdot 10^{-4}$ | 204, 920 | odds ratio: 2.943 |
| line 107 | chi-square test | prefrontal readers vs non-readers likeliness to participate in candidate cell assemblies targeting amygdalar readers | $p = 2.6 \cdot 10^{-21}$ | 404, 2018 | odds ratio: 1.797 |
| 3.a center | Sign test, one-tailed | reader response vs proportional response to co-activation of up to one third of members | $p = 0.7186$ | 967 | $z = -0.579$ |
| 3.a center | Sign test, one-tailed | reader response vs proportional response to co-activation of more than one third and up to two thirds of members | $p = 1.3748 \cdot 10^{-13}$ | 1247 | $z = 7.306$ |
| 3.a center | Sign test, one-tailed | reader response vs proportional response to co-activation of more than two thirds of members (but not all members) | $p = 0.7186$ | 613 | $z = 3.312$ |
| 3.a bottom | Wilcoxon signed-rank test | linear vs sigmoidal fit | $p = 2.844 \cdot 10^{-132}$ | 1026 | $z = -24.473$ |
| 3.b bottom | Wilcoxon rank sum test | sparsity increase non-readers vs readers | $p = 2.5792 \cdot 10^{-12}$ | 2018, 404 | $z = -6.999$ |
| 3.c | Wilcoxon rank sum test | Discrimination indices observed vs shuffled data | $p = 3.9281 \cdot 10^{-27}$ | 100, 10000 | $z = 10.788$ |
| 4.b | chi-square test | proportion of AMY-PFC assembly–reader pairs significantly changed their responses following fear conditioning vs following control sessions | $p = 0.0017$ | 171, 260 | odds ratio: 1.677 |
| 4.b | chi-square test | proportion of AMY-PFC assembly–reader pairs significantly changed their responses following fear extinction vs following control sessions | $p = 0.7253$ | 235, 260 | odds ratio: 1.106 |
| 4.b | chi-square test | proportion of PFC-AMY assembly–reader pairs significantly changed their responses following fear conditioning vs following control sessions | $p = 0.1403$ | 171, 267 | odds ratio: 1.582 |
| 4.d | chi-square test | proportion of PFC-AMY assembly–reader pairs significantly changed their responses following fear extinction vs following control sessions | $p = 0.0347$ | 123, 267 | odds ratio: 2.060 |

